# maGENEgerZ: An Efficient AI-Based Framework Can Extract More Expressed Genes and Biological Insights Underlying Breast Cancer Drug Response Mechanism

**DOI:** 10.1101/2023.12.29.573686

**Authors:** Turki Turki, Y-h. Taguchi

**Affiliations:** King Abdulaziz University, Department of Computer Science, Jeddah 21589, Saudi Arabia.; Chuo University, Department of Physics, Tokyo 112-8551, Japan.

**Author notes:** **Corresponding Authors**. Turki Turki and Y-h. Taguchi.

**Keywords:** Breast cancer, drug response, gene expression, machine learning, deep learning, AI application in cancer clinical trials

## Abstract

Understanding breast cancer drug response mechanism can play a crucial role in improving the treatment outcomes and survival rates. Existing bioinformatics-based approaches are far from perfect and do not adopt computational methods based on advanced artificial intelligence concepts. Therefore, we introduce a novel computational framework based on an efficient support vector machines (esvm) working as follows. First, we downloaded and processed three gene expression datasets related to breast cancer responding and non-responding to the treatments from the gene expression omnibus (GEO) according to the following GEO accession numbers: GSE130787, GSE140494, and GSE196093. Our method esvm is formulated as a constrained optimization problem in the dual form as a function of λ. We recover the importance of each gene as a function of λ, y, and x. Then, we select *p* genes out of *n,* provided as input to enrichment analysis tools, Enrichr and Metascape. Compared to existing baseline methods including deep learning, results demonstrate superiority and efficiency of esvm achieving high performance results and having more expressed genes in well-established breast cancer cell lines including MD-MB231, MCF7, and HS578T. Moreover, esvm is able to identify (1) various drugs including clinically approved ones (e.g., tamoxifen and erlotinib); (2) seventy-four unique genes (including tumor suppression genes such as TP53 and BRCA1); and (3) thirty-six unique TFs (including SP1 and RELA). These results have been reported to be linked to breast cancer drug response mechanism, progression, and metastasizing. Our method is available publicly in the maGENEgerZ web server at https://aibio.shinyapps.io/maGENEgerZ/.

## 1. Introduction

The ability to elucidate various mechanisms underlying breast cancer drug response and resistance is a critical part in the clinical decision-making process, not only aiding in finding out the potential effectiveness of a drug compound but also spanning to (1) reducing the search space for candidate compounds; (2) having a greater awareness and management of probable adverse reactions before conducting clinical trials [1]; and (3) identifying potential drug targets associated with drug compounds. Studies have been conducted to analyze gene expression data obtained from biological experiments pertaining to breast cancer drug response to unveil various molecular mechanisms. Du et al. [2] utilized a bioinformatics approach to identify important genes that play a key role in overcoming breast cancer drug resistance working as follows. First, two gene expression datasets were downloaded from the gene expression omnibus (GEO) database based on GEO accession numbers GSE28694 (Miller & Payne grades 4 and 5) and GSE28826 (Miller & Payne grades 1 and 2). The GSE28694 dataset had 13 samples treated as drug-sensitive group while the GSE28826 dataset had 28 samples treated as drug-resistant group. Both processed datasets were provided as input to limma to identify 255 differentially expressed genes (DEGs) with *p* < 0.05, assigned to enrichment analysis tool ClusterProfiler. The protein-protein interaction with the use of random walk identified three genes (i.e., PRC1, GGTLC1, and IRS1) that are involved in immune pathways and involved in the breast cancer drug resistance mechanism. Further validation of the importance of these three genes was performed using additional datasets from the GEO and TCGA databases.

Wu et al. [3] utilized a bioinformatics approach to identify a gene signature that aid in predicting neoadjuvant chemotherapy response for breast cancer patients. They downloaded a gene expression dataset from the GEO database according to the GEO accession number GSE25066. The processed dataset had 508 samples in which 16 samples were excluded because of missing data, resulting in 492 samples. To perform differential gene expression analysis, limma was applied to drug-resistant tumor samples against drug-sensitive tumor samples identifying 347 DEGs. Then, applying limma within the drug-resistant cell line samples against wild type cell samples to identify 296 DEGs. Then, identifying 36 genes common between the 347 and 296 DEGs. The 36 genes were provided as input to enrichment analysis finding out 12 hub genes considered as a gene-signature (HJURP, IFI27, RAD51AP1, EZH2, DNMT3B, SLC7A5, DBF4, USP18, ELOVL5, PTGER3, KIAA1324, and CYBRD1) from the PPI that has been validated to assess its discriminative power using lasso in which the same GSE25066 dataset was divided into training and validation sets.

Freitas et al. [4] performed a bioinformatics analysis to identify reliable biomarkers for adding carboplatin to the standard anthracycline/taxane treatment, which can aid in identifying triple negative breast cancer (TNBC) patients achieving pathological complete response to neoadjuvant chemotherapy (NAC). Therefore, TNBC patients with expected poor clinical outcomes can be provided another treatment options. The processed gene expression data was for 66 patients in which 33 were treated with carboplatin + paclitaxel (composed of 19 having RD and 14 achieving PCR) while the remaining 33 were treated with paclitaxel (composed of 23 having RD and 10 achieving PCR). In the 33 patients treated with carboplatin + paclitaxel, they applied limma to identify 37 DEGs between RD and PCR while identifying 27 DEGs between RD and PCR patients treated only with paclitaxel. Moreover, identifying 24 DEGs between RD and PCR patients for the 66 patients. Then, selecting 10 statistically significant genes (BNIP3, ZBTB16, KCNB1, HAS1, HEMK1, TFF1, PLA2G4F, SNAI1, C5orf38 and GRIN2A) out of the 37 and 27 DEGs, and 3 statistically significant genes (ALDH1A1, MCM2, and CXADR) out of the 27 DEGs. These 13 genes acted as gene signatures and reported results demonstrated their feasibility to discriminate between patients having RD and those achieving PCR.

Stevens et al. [5] aimed to unveil the molecular mechanism behind inducing a chemotherapy-resistant in inflammatory breast cancer (IBC) patients. They had a dataset of 131 samples between IBC and non-IBC patients derived from several profiling methods distributed based on 14 samples using ChIP-seq profiling, 84 samples using RNA-seq profiling, 3 samples using single-cell RNA-seq profiling, and 30 samples using RNA-seq II profiling. The dataset was deposited into the GEO database with accession number GSE163397. Bioinformatics analysis coupled with enrichment analysis revealed that JAK2/STAT3 signaling is a key player in driving a chemo-resistance in IBC. Therefore, inhibition of JAK2/STAT3 coupled with the use of paclitaxel drug can overcome therapeutic resistance in IBC patients. Debets et al. [6] performed a bioinformatics analysis coupled with enrichment analysis to identify a molecular signature (ER2, HER4, ER, IGF1R, and Kalirin) predictive of treatment response and resistance in HER2-positive breast cancer patients. Miri et al. [7] performed a bioinformatics analysis to identify critical genes and pathways that play a key role in doxorubicin resistance in breast cancer. They downloaded two gene expression datasets from GEO with GSE24460 and GSE76540 accession numbers. Then, applying limma to normal and resistant samples to doxorubicin, identifying 1,108 and 3,207 DEGs in GSE24460 and GSE76540, respectively. Pearson correlation was performed to select 36 and 406 significant genes in GSE24460 and GSE76540. Gene co-expression network (GCN) analysis was performed to identify 18 and 115 genes in GSE24460 and GSE76540, respectively. Nine genes (ABCB1, MMP1, TCEAL2, AKAP12, PLS3, LDHB, NEFH, CNN3, and VIM) were common between the two datasets, reported to play a key role in doxorubicin resistance. Other studies aimed to unveil various mechanisms leading to drug resistance in breast cancer patients [8–13].

Although current advancing in breast cancer drug mechanism within clinical testing mainly depends on bioinformatics-driven computational approaches with existing off-the-shelf tools, AI-driven computational frameworks are needed to unveil vast biological insights and to properly promote the use of AI in real clinical settings. The availability of such AI tools can help clinical oncologists to early avoid therapeutic targets associated with poor treatment responses and thereby advancing clinical understanding and reducing the search space for potential drugs with adverse effects. The novelty in our study is ascribed to the following major contributions:

(1) We introduce an AI-driven computational approach consisting of an efficient support vector machines (esvm) combined with enrichment analysis tools (Enrichr and Metascape), unveiling various molecular mechanisms pertaining to breast cancer drug response [14–16].
(2) We downloaded and processed three gene expression datasets pertaining to breast cancer drug response according to the following GEO accession numbers: GSE130787, GSE140494, and GSE196093.
(3) Performing an extensive experimental study from biological and classification perspectives, comparing our method against other bioinformatics-based methods (limma [17], sam, *t*-test [18, 19], and lasso [20]) and adapted deep learning methods (DeepLIFT [21] , DeepSHAP [22], and LRP [23]).
(4) Compared to all methods including deep learning-based methods, experimental results based on enrichment analysis demonstrate that our method (esvm) identified more expressed genes in three well-established breast cancer cell lines, including MD-MB231, MCF7, and HS578T. Moreover, we identified various drugs for breast cancer including FDA-approved ones such as Gemcitabine (Gemzar) and tamoxifen (Nolvadex). Moreover, identifying 74 unique genes including tumor suppression genes such as TP53, PTEN, BRCA1, and RB1. Reporting 36 unique TFs including SP1, NFKB1, and RELA. All of which have been reported to play a key role in breast cancer drug response and resistance mechanism. In terms of the running time for learning-based methods, lasso was faster than our method esvm in which both were computationally faster than all other deep learning-based methods.
(5) Results from a classification perspective demonstrated the superiority of genes set obtained via our method when coupled with learning algorithms. Specifically, In Dataset1 when balanced accuracy (BAC) is considered, SVM coupled with a gene set from our method achieved a 32.4% performance improvement over the second-best for SVM with all genes (named None) (see Table S4 in Supplementary Additional file). In Dataset2 using BAC performance measure (see Table S5 in Supplementary Additional file), ours when coupled with SVM had 38.1% performance improvement when compared to the second-best for SVM coupled with gene set from sam. In the last dataset (i.e., Dataset3) as shown in Table S5 of Supplementary Additional file, SVM coupled with gene set from our method had a 6.1% performance improvement over the second-best for SVM coupled with gene set from DeepLIFT. The same holds true when we evaluated the classification performance using lasso as a learning algorithm coupled with gene set from our method.
(6) For reproducibility of the analysis in this study, we made a publicly available implementation for our method esvm within the maGENEgerZ web server at https://aibio.shinyapps.io/maGENEgerZ/. Moreover, we included the processed datasets within the Supplementary Datasets folder. We also provided a Supplementary maGENEgerZ_Screenshots.docx file to show the use for our web tool.

## 2. Materials and Methods

### 2.1 Gene Expression Profiles

In this work, we retrieved three datasets from different gene expression experiments of different GEO accession numbers [24].

#### 2.1.1 GSE130787: Dataset1

For this dataset derived from the gene expression experiment at the GEO database, we had 89 samples and 5267 genes. As a result, we encoded the dataset as an 89 × 5268 matrix including drug responses as a column vector. The 89 samples were distributed in terms of treatment into three groups as follows. Twenty-six samples for patients treated with docetaxel, carboplatin, and trastuzumab (TCH). Thirty-eight samples for patients treated with docetaxel, carboplatin, trastuzumab and lapatinib (TCHTy). Twenty-five samples for patients with docetaxel, carboplatin and lapatinib (TCTy). In terms of the distribution of drug responses, thirty-eight BC patients achieved pathological complete response (PCR) while fifty-one BC achieved residual disease (RD). The gene expression experiment was performed using the microarray platform, Agilent-014850 Whole Human Genome Microarray 4×44K G4112F (Probe Name version). This dataset is referred to as Dataset1.

#### 2.1.2 GSE140494: Dataset2

For this second dataset derived from the performed gene expression experiment, We had 91 samples and 5313 genes, which were approved as protein-coding genes (PCGs) by domain experts from HUGO Gene Nomenclature Committee (HGNC) Biomart at https://biomart.genenames.org/ [25, 26]. Thus, we encoded the dataset as a 91 × 5314 matrix, including a column vector for drug responses. The drug responses were distributed as follows. Nineteen BC patients achieved resistant response while seventy-two were sensitive to the treatment (i.e., docetaxel, followed by 5-fluorouracil, epirubicin, and cyclophosphamide (TFEC). The gene expression experiment was performed using the microarray platform, Affymetrix Human Genome U133 Plus 2.0 Array. This dataset is referred to as Dataset2.

#### 2.1.3 GSE196093: Dataset3

For this third dataset, we had 736 samples and 118 genes. Consequently, we encoded the dataset as a 736 × 119 matrix, including a column vector for drug responses in which 256 BC patients achieved complete response (CR) while 480 had a failed complete response (FCR). We had 11 treatments in which the 736 samples were distributed accodingly as follows. Paclitaxel (169), Paclitaxel + ABT 888 + Carboplatin (63), Paclitaxel + AMG-386 (110), Paclitaxel + AMG-386 + Trastuzumab (18), Paclitaxel + MK-2206 (56), Paclitaxel + MK-2206 + Trastuzumab (31), Paclitaxel + Neratinib (105), Paclitaxel + Pembrolizumab (67), Paclitaxel + Pertuzumab + Trastuzumab (43), Paclitaxel + Trastuzumab (25), and T-DM1 + Pertuzumab (49). The gene expression data was performed using Reverse Phase Protein Array (RPPA) microarray at George Mason University. This dataset is referred to as Dataset3. Table 1 provides an overview for the three-studied datasets.

**Table 1:** Overview of the three studied breast cancer drug response datasets downloaded from the gene expression omnibus database.

| Dataset | Samples | Responder | Non-responder | Genes | Platform | Organism | Experiment Type |
| --- | --- | --- | --- | --- | --- | --- | --- |
| GSE130787 | 89 | 38 | 51 | 5267 | GPL6480 | Homo sapiens | Expression profiling by array |
| GSE140494 | 91 | 72 | 19 | 5313 | GPL570 | Homo sapiens | Expression profiling by array |
| GSE196093 | 736 | 256 | 480 | 118 | GPL28470 | Homo sapiens | Protein profiling by protein array |

### 2.2 Computational Framework

In Figure 1, we outline the main steps pertaining to our computational framework. In terms of the preprocessing part, biopsy samples were obtained from breast cancer patients. Then, collected samples were prepared and provided to a biological technology, measuring the gene expression levels [27]. In the machine learning part, the input data corresponds to a gene expression dataset, where x*_i_* represents the *i*th sample and y*_i_* is the associated drug response. In our study, y*_i_* is a binary class label (e.g., {pathological complete response (PCR), residual disease (RD)}). The entire samples x*_i_* (for *i =* 1..*m*) were encoded as an *m* × *n* matrix in which *m* and *n* are the number of samples and genes, respectively. All drug responses y*_i_* (for *i =* 1..*m*) encoded as a 1 × *m* column vector. To identify *p* important genes out of the *n* genes in which *p* ≪ 𝑛, we used to find arguments (i.e., w = [*w*_1_…*w*_n_] and *b* ∊ R) that minimize the objective function in Equation 1 subject to the linear constraints as in [28, 29]. After solving the optimization problem in Equation 1, weights in w correspond to the importance of genes in which higher weights indicate the more important these genes are. However, the main issue is as follows.

**Figure 1:**
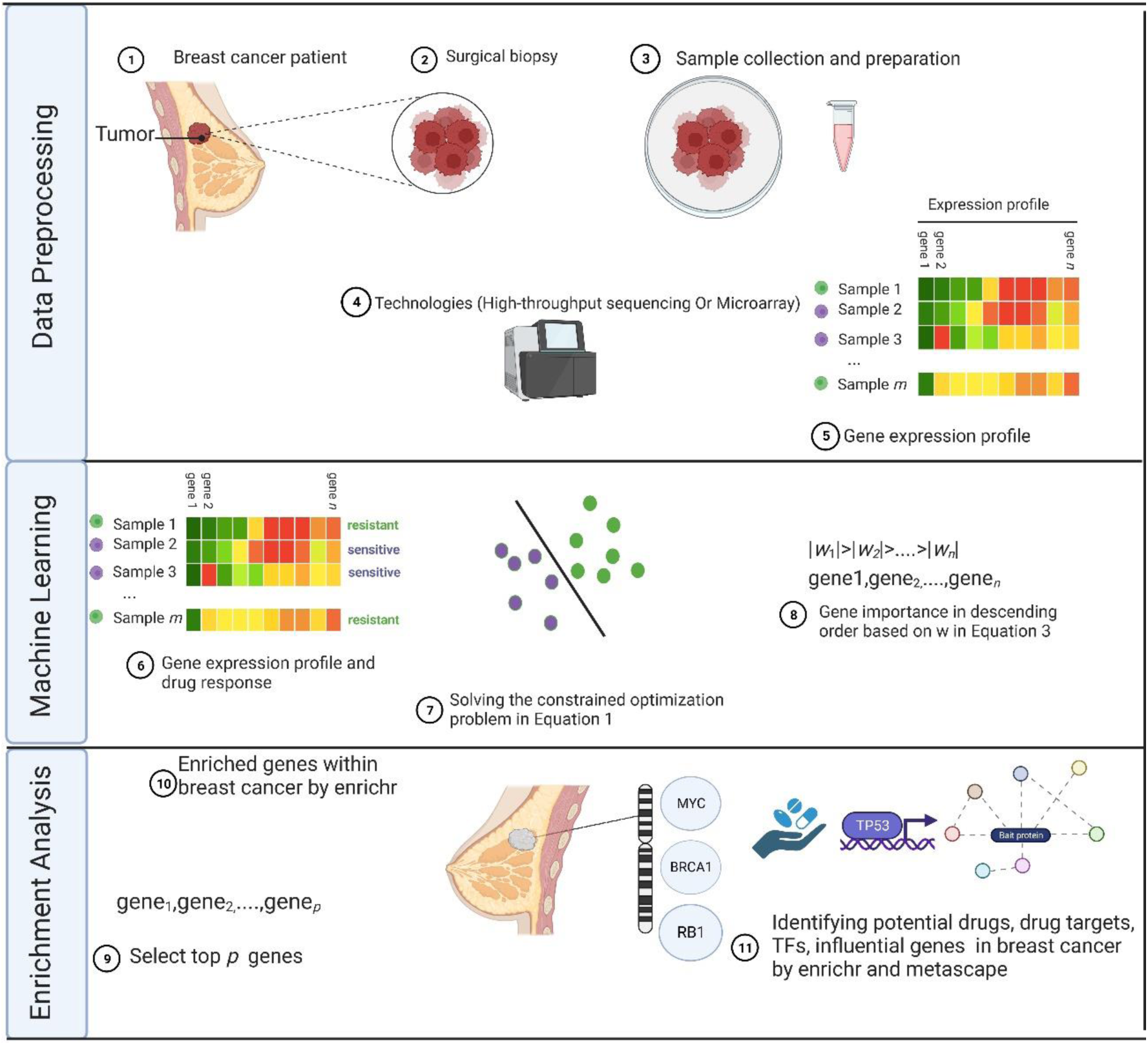
Flowchart of the introduced AI-based framework identifying drugs, drug targets, critical genes, and transcription factors in breast cancer. Figure created with BioRender.com.

The optimization problem in Equation 1 depends on w and *b,* where |W| *is* equal to *n,* which is way larger than *m* (i.e., 𝑚 ≪ 𝑛) in genomic sciences [30]. That makes the solution for the optimization problem in Equation 1 computationally expensive [31, 32], where the number of genes (*n*) is typically larger than the number of samples (*m*).

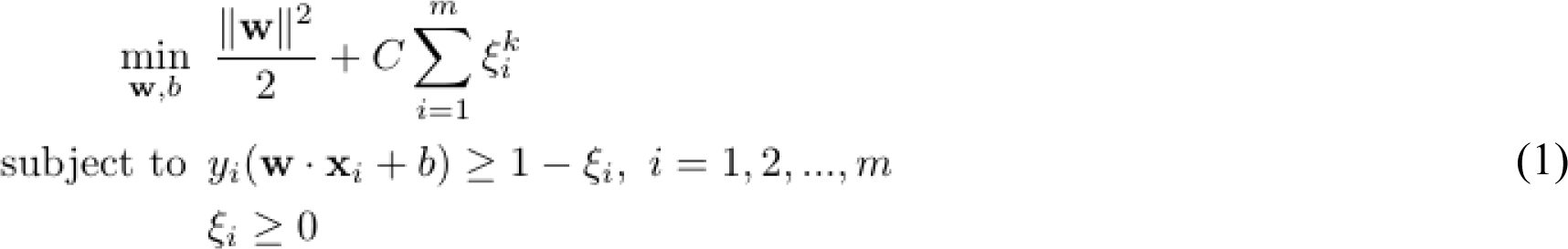

Therefore, we seek to solve the dual form of SVM (See Equation 2)[31]. It can be seen that the optimization problem now depends on finding the Lagrange multiplier (λ) in which each 𝑥_𝑖_ is associated with λ_𝑖_. This is way faster than finding w in Equation 1[31].

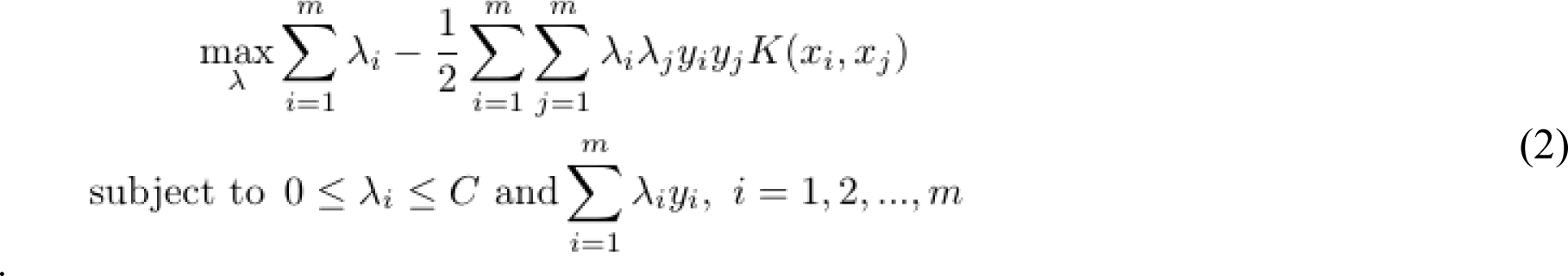

In Equation 3 [31], we recover w from λ in Equation 2 as follows:

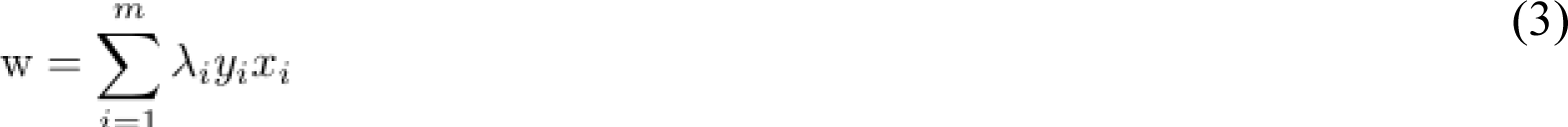

For any 𝑥_𝑖_ associated with 0 < λ_𝑗_ < 𝐶, *b* is recovered as

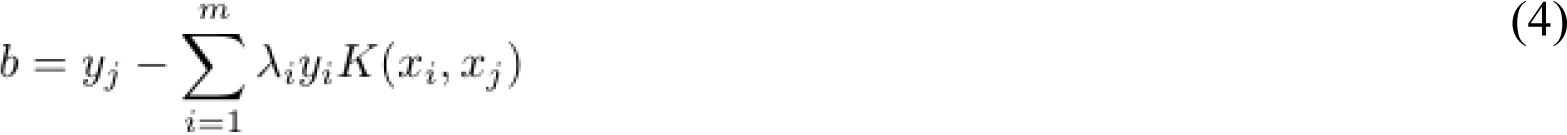

where 𝐾(𝑥_𝑖_, 𝑥_𝑗_) is a similarity measure *K* (usually called kernel) of 𝑥_𝑖_ and 𝑥_𝑗_.

A testing example 𝑧 is predicted as

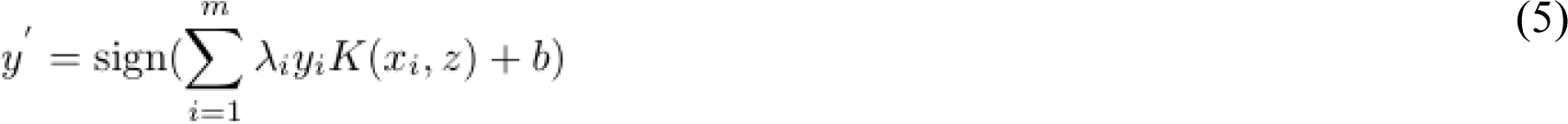

where sign() is an indicator function mapping to 1 (corresponding to RD) if its argument is greater than or equal to 0. Otherwise, it is mapped to -1 (corresponding to PCR). Eqs. S1-S5 in Supplementary Additional file show 5 popular kernels used with SVM [31]. When a linear kernel is used, the prediction model becomes as

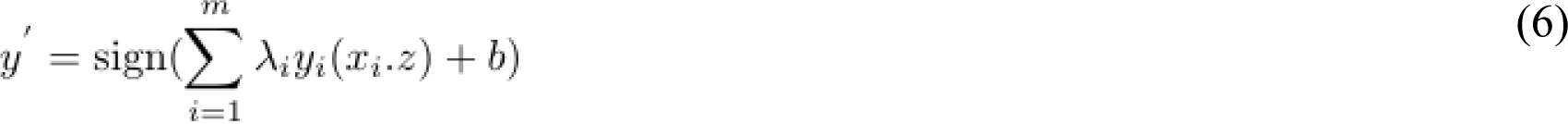

We used CVXR in R to solve the dual form of SVM in Equation 2 and to find λ [33]. In terms of the enrichment analysis part, we uploaded the *p* genes as input to Enricher and Metascape, where these *p* genes are weighted with the top *p* weights (|*w*_1_|>|*w*_2_|>…>|*w_p_*|). Then, interpreting and identifying biologically related terms. Including key expressed genes, drugs, drug targets, transcription factors, and others are provided in the next section.

## 3 Experiments and Results

### 3.1 Experimental Methodology

We compared our method esvm against the following baseline methods: linear models for microarray data (limma) [17], significance analysis of microarrays (SAM), Student’s t-test (*t*-test), and least absolute shrinkage and selection operator (lasso) [18]. The input to the 5 studied methods is a labeled gene expression data. Because the prediction in our efficient svm-based model (named esvm) is defined as the sign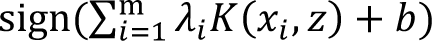, we had to recover w as 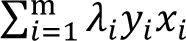 and then selecting *p* genes associated with the top *p* weights (i.e., w = [|*w*_1_|…|*w_p_*|]), corresponding to the top *p* important genes. For lasso, as the model is expressed as 𝛽_0_ + 𝛽x, we selected the *p* genes associated with top *p* coefficients (i.e., 𝛽 = [|𝛽_1_|…|𝛽*_p_*|]) (excluding genes associated with zero coefficients). For limma, SAM, and *t*-test baseline methods, genes were selected based on according to significantly adjusted *p*-values < 0.01.

To perform enrichment analysis and evaluate the results from biological perspective, we uploaded genes obtained from each method to Enrichr (https://maayanlab.cloud/Enrichr/) and Metascape (https://metascape.org/gp/index.html) [14, 16]. When retrieved terms are related to breast cancer, then a method that has terms associated with more genes is considered the superior method. Furthermore, we assessed the performance from the classification perspective against lasso as a baseline, reporting area under ROC curve (AUC) as a performance measure, followed by conducting an statistical significance test and reporting the running time. In this study, we utilized R to run the experiments [34]. Specifically, we used the CVXR package in R to aid in solving the formulated optimization problem [33]. We employed siggenes package to run SAM [35] and we employed the limma package in R to run limma using the the two functions, lmFit and eBayes [17]. In terms of the *t*-test, we employed the t.test function within the stats package [34]. To run lasso, we used the glmnet package [20] in which we set λ=0.05 and also utilized cross-validation on the training set to find the optimal λ when using lasso for classification. For SAM, limma, and *t*-test, to compute adjusted *p*-values, we employed the p.adjust function with the “BH”, setting *p* < 0.01 as in [36, 37]. We used DeepLift, DeepSHAP, and LRP functions in innsight package in R to run DeepLIFT, DeepSHAP, and LRP, respectively [38].

For Dataset1, the gene expression experiment is to analyse and evaluate neoadjuvant docetaxel and carboplatin plus trastuzumab and/or lapatinib for patients with HER2+ breast cancer, obtained from the school of medicine at department at the University of California, Los Angeles, USA. We retrieved gene expression profiles of patients responding to the treatment (labelled as pathological complete response (PCR)) and those not completely responding to the treatment (labelled as residual disease (RD)) from the gene expression omnibus (GEO) database (https://www.ncbi.nlm.nih.gov/geo/) with GEO accession number GSE130787. We utilized getGEO() function within GEOquery package [39] to download and get the gene expression data. We utilized the fData() function from the Biobase package [40] to process and get gene names, pData() function within the Biobase package was used to get drug responses (i.e., PCR and RD) associated with each sample, exprs() function within the Biobase package to get expression values, missForest() function within missForest package to impute missing values [41, 42]. We selected 5267 protein coding genes (PCGs) according to Biomart domain experts at HUGO Gene Nomenclature Committee (See subsection GSE130787: Dataset1).

For Dataset2, the gene expression experiment is for predicting neodjuvant chemotherapy response in early breast cancer, obtained from leibniz research centre for working environment and human factors, located at Dortmund, Germany. We retrieved gene expression profiles of breast cancer patients responding (labelled as PCR and pathological partial response (PPR)) and those not responding to treatment (labelled as pathological no change (PNC)) from the GEO database NIH under GEO accession number GSE140494. As in Dataset1, we used the five functions (i.e getGEO(), fData(), pData(), exprs(), and missForest()) to download, prepare and impute Dataset2. Moreover, we selected 5313 PCGs according to BioMart domain experts at HUGO Gene Nomenclature Committee (See subsection GSE140494: Dataset2).

For Dataset3, the gene expression experiment is for I-SPY 2 neoadjuvant chemotherapy/targeted-therapy trial for early-stage breast cancer patients with high risk, obtained from University of California, San Francisco, USA. Gene expression profiles pertaining to breast cancer patients responding to treatment (labelled as complete response) against those not responding properly to the treatment (labelled as failed complete response) were obtained from the GEO database under GEO Accession number GSE196093. As in Dataset2, we used the five functions (i.e getGEO(), fData(), pData(), exprs(), and missForest()) to download, prepare and impute Dataset3.

### 3.2 Results

#### 3.2.1 Dataset1

In Table 2, we report terms obtained from Enrichr based on input genes provided via each method. Terms in Table 2(a) show expressed genes within breast tissues according to each method. The more expressed genes, the better the computational method is. Table 2(a) demonstrates that esvm outperformed all compared methods, obtaining 28 expressed genes (SLC22A3, FKBP10, GRP, STAB2, LPL, GLI3, ADCY5, OBP2A, SOSTDC1, APOD, HMGCS2, GABRE, CCL21, TMEM61, BBOX1, GFRA1, IGF1, BMP6, APLN, CAPN13, NR4A1, C1ORF116, ANGPTL7, KCNS3, SSPN, PGR, FGFR2, and LTF) within the breast tissue. The second-best method is limma with 14 expressed genes (TMEM86B, LRP1, MEOX1, PLCZ1, OBP2A, PPP1R1B, LRIG1, ECHDC3, MAFK, MZF1, A2M, RNF186, FMOD, and KANK3) within the breast tissue. The worst-performing method is lasso in which 8 expressed genes (DGAT1, RNASE7, FXYD1, VEGFB, ROR2, FGF1, HSPA12A, and PTPN14) are expressed in the breast tissue. In Table 2(b), we show retrieved terms (i.e., cancer cell lines) and expressed genes within NCI-60 Cancer Cell Lines. Our method, esvm, performed better than other methods, obtaining a total of 8 expressed genes within the retrieved breast cancer cell lines. Specifically, three genes (PNMT, COL13A1, BCAS1) were expressed within MD-MB231, three genes (MYO5C, GFRA1, FGFR2) were expressed within MCF7, and two genes (COL13A1 and FAR2) were expressed within HS578T. The second-best method is limma (tied with sam), both having 4 expressed genes within two breast cancer cell lines. In terms of limma, one gene (ECHDC3) was expressed within MD-MB231, whereas three genes (RBM8A, USP18 and PI4K2A) were expressed within MCF7. For sam, one gene (CLSPN) was expressed within MD-MB231, three genes (TFAP2C, GFRA1, and CNNM3) were expressed within MCF7. The worst-performing method is lasso, having two expressed genes within breast cancer cell lines. These results demonstrate the superiority for our method (i.e., esvm) in identifying breast tissue as well as breast cancer cell lines used in cancer research. In Supplementary Table1_A and Table1_B, we include enrichment analysis results of ARCHS4 Tissues and NCI-60 Cancer Cell Lines, respectively.

**Table 2:**
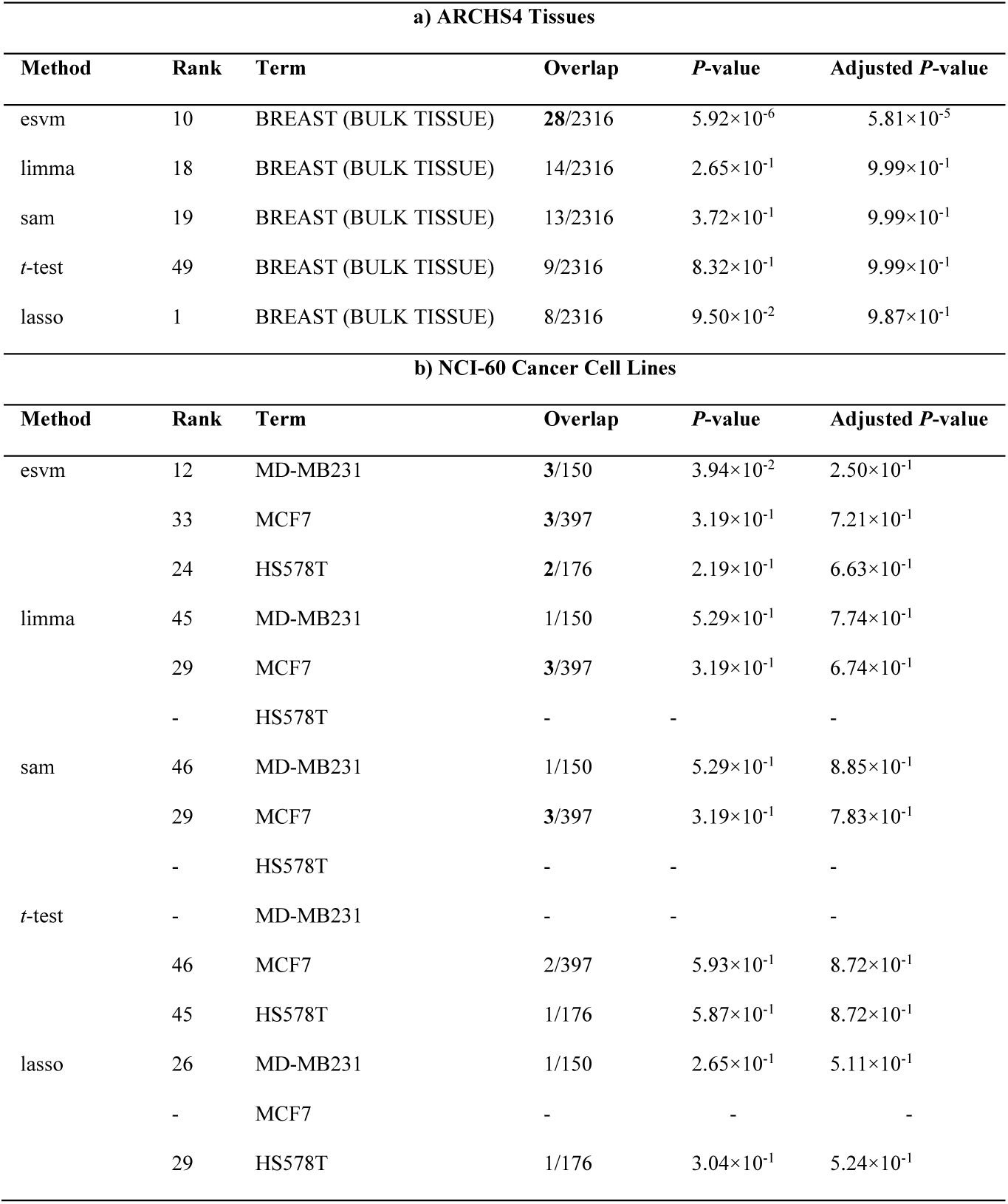
Enriched terms obtained from a) ARCHS4 Tissues and b) NCI-60 Cancer Cell Lines via Enrichr according to produced genes by each method when using Dataset1. The best result is shown in bold.

In Figure 2(a), we provide a visualization of intersected genes produced by all computational methods. It can be seen that esvm has unique 94 genes out of 100 when compared to all other methods. This implies that each method generated a different list of genes. For example, limma, sam, *t-*test, and lasso had unique genes of 96, 95, 94, and 38, respectively. It can also be seen that the number of common genes between any pair of methods is at most 2. This indicates that each method generated a different gene set and similarity among methods is minimal. In Supplementary DataSheet1_B, we list the genes according to the UpSet plot in Figure 2(a).

**Figure 2:**
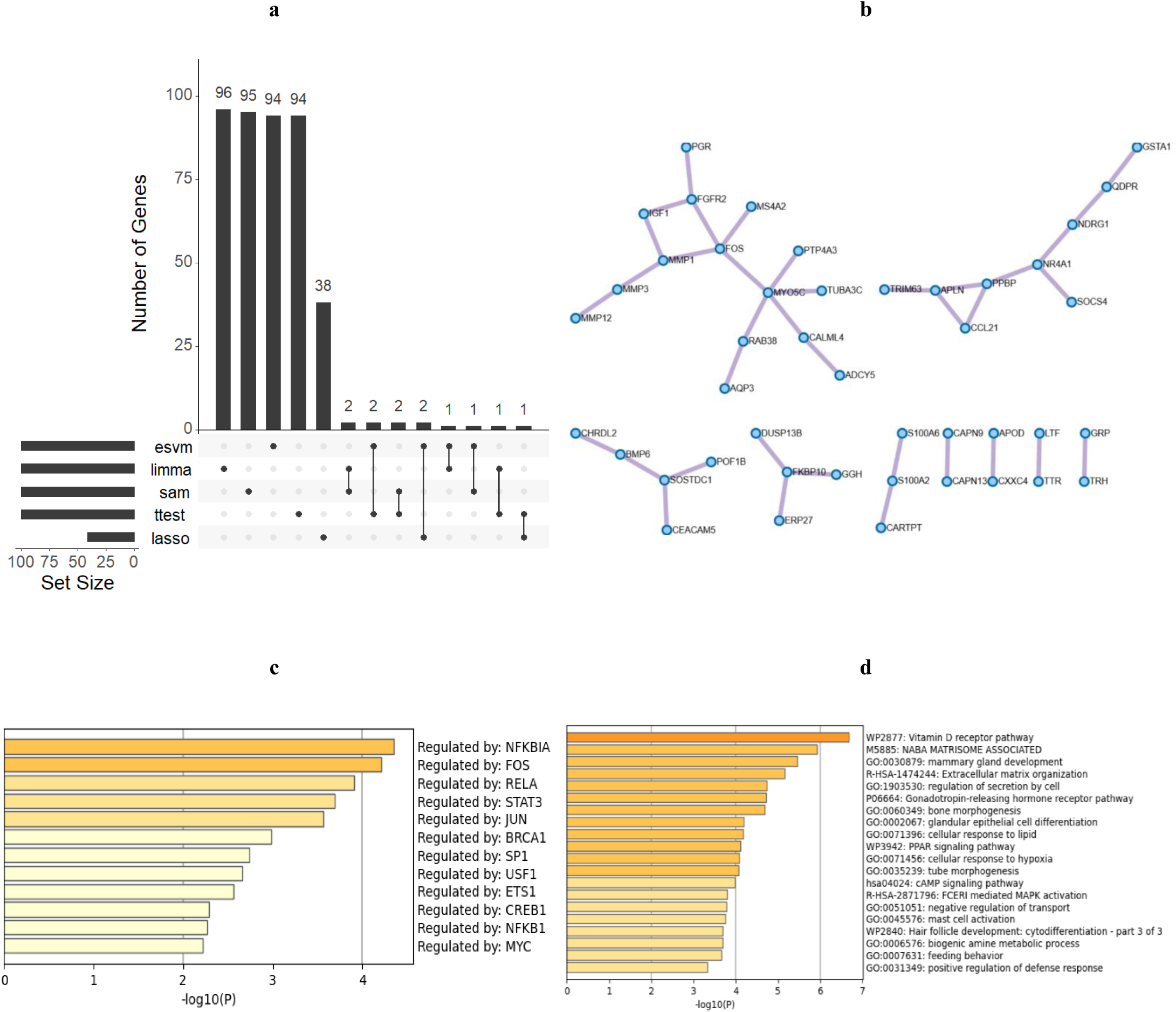
(a) UpSet plot of gene lists provided by the computational methods when using Dataset1. (b) Nine clusters in a protein-protein interaction based on genes of esvm coupled with Metascape. (c) Twelve transcription factors according to Metascape when coupled with genes from esvm. (d) Process and pathway enrichment analysis provided by Metascape according to the 100 genes of esvm.

As esvm demonstrated superiority among the other computational methods, we provide the 100 genes produced via esvm to Metascape for further enrichment analysis. Figure 2(b) displays the following 44 genes obtained from protein-protein interaction: GRP, TRH, LTF, TTR, APOD, CSSC4, CAPN9, CAPN13, S100A6, S100A2, CARTPT, DUSP13B, FKBP10, GGH, ERP27, POF1B, SOSTDC1, CEACAM5, BMP6, CHRDL2, GSTA1, QDPR, NDRG1, NR4A1, SOCs4, PPBP, CCL21, APLN, TRIM63, ADCY5, CALML4, MYO5C, RAB38, AQP3, TUBA3C, PTP4A3, FOS, MS4A2, MMP1, MMP3, MMP12, FGFR2, IGF1, and PGR. The enrichment analysis showed that 10 genes (BMP6, IGF1, MMP1, MMP3, MMP12, PPBP, S100A2, S100A6, CCL21, CHRDL2) were related to NABA MATRISOME ASSOCIATED, 7 genes (BMP6, IGF1, PPBP, S100A2, S100A6, CCL21, CHRDL2) were related to NABA SECRETED FACTORS, and 6 genes (BMP6, FGFR2, NR4A1, IGF1, MMP12, APLN) were related to positive regulation of epithelial cell proliferation. It has been reported that NABA MATRISOME ASSOCIATED and NABA SECRETED FACTORS pathways play a key role in breast cancer metastasis through its involvement with extracellular matrix proteins [43, 44]. Also, genes linked to positive regulation of epithelial cell proliferation biological process are related to induction of metastasis and inhibition of breast cancer cells apoptosis through the promotion of epithelial cell proliferation via estrogen [45, 46]. These results demonstrate the effectiveness of our computational method in unveiling important molecular mechanisms pertaining to breast cancer pathogenesis and metastasis.

Figure 2(c) reports 12 transcription factors (TFs): NFKBIA, FOS, RELA, STAT3, JUN, BRCA1, SP1, USF1, ETS1, CREB1, NFKB1, and MYC. BRCA1 is known to play a key role in various biological processes in breast cancer [47–51]. TFs such as ETS1 and STAT3 have been reported as potential therapeutic targets in breast cancer [52] . Suppression of NFKBIA and CREB1 have been reported to be related to the inhibition of breast cancer progression [53, 54] . These 12 TFs can (1) aid in understanding breast cancer molecular mechanism; and (2) act as potential therapeutic targets for breast cancer treatment. In Figure 2(d), genes provided via esvm are related to various biological processes and pathways in breast cancer progression. The top enriched term is vitamin D receptor pathway, which ameliorates breast cancer via contributing to the growth regulation of breast cancer cells [55, 56]. Other enriched terms such as NABA MATRISOME ASSOCIATED, mammary gland development, and extracellular matrix organization have been reported to play a key role in various biological processes to breast cancer progression and treatment [57–61]. In Supplementary Enirchment_Dataset1, we list Metascape enrichment analysis results related to Dataset1.

In Table 3, we report terms (i.e., drugs) and expressed genes (i.e., drug targets) within IDG Drug Targets 2022. Tamoxifen and Fulvestrant are antiestrogen inhibitors (hormone therapy) that have been approved for breast cancer treatment (See Figure 3) [62–64]. Both drugs identified two drug targets, PGR, ATP1A2. Cisplatin is a chemotherapy used for breast cancer treatment [65, 66]. ATP1A2 was a drug target reported in association with cisplatin. The obtained biological knowledge contributes to more understanding of breast cancer progression and treatment. In Supplementary Table1_C, we provide enrichment analysis results pertaining to IDG Drug Targets 2022.

**Table 3:** Retrieved enriched terms from IDG Drug Targets 2022 via Enrichr according to uploaded genes from esvm.

| Rank | Term | Class | Genes |
| --- | --- | --- | --- |
| 35 | Tamoxifen | Antiestrogen | PGR;ATP1A2 |
| 37 | Fulvestrant | Estrogen Receptor Antagonist | PGR |
| 64 | Cisplatin | Platinum Coordination Complex | ATP1A2 |

**Figure 3:**
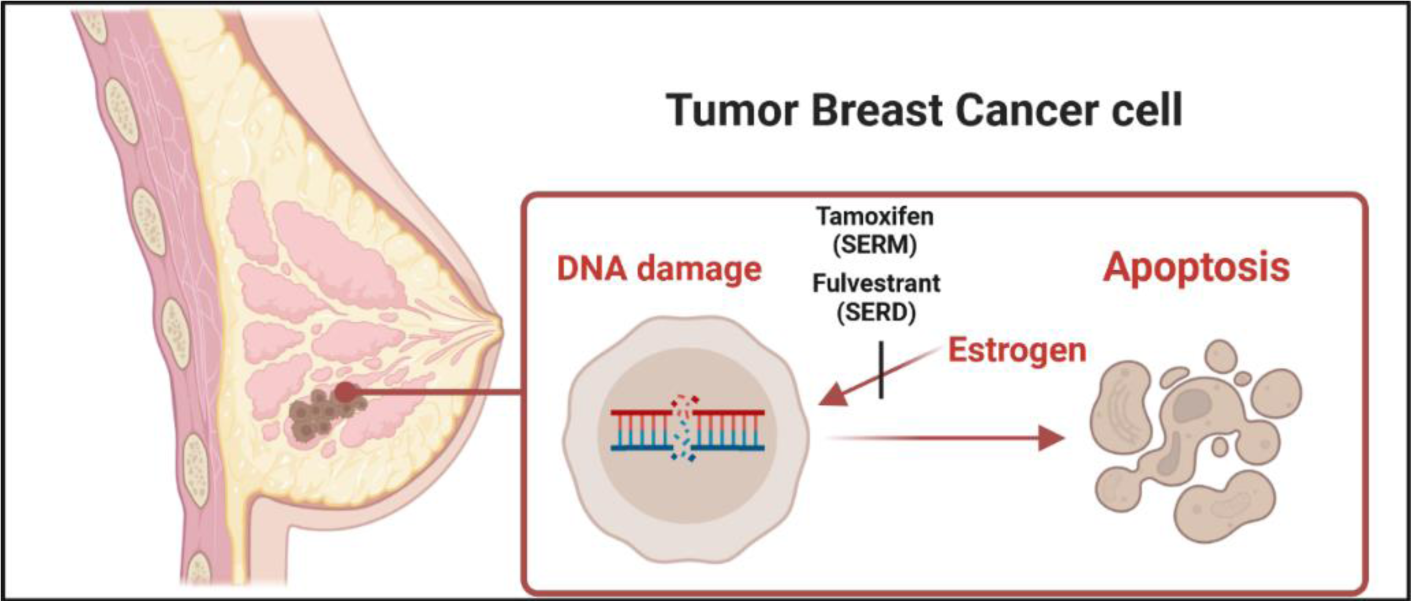
Tamoxifen and fulvestrant anticancer drugs inhibiting estrogen, causing DNA damage and cell death. SERM is selective estrogen receptor modulator. SERD is selective estrogen degrader.

#### 3.2.2 Dataset2

In Table 4(a), we report retrieved breast tissue terms associated with expressed genes while Table 4(b) reports breast cancer cell line terms with expressed genes after uploading genes produced via all computational methods. It can be seen from Table 4(a) that our method (esvm) performed better than its competing baseline methods. Particularly, esvm had breast tissue term with 20 expressed genes (COL15A1, PFKFB3, VCAM1, TAT, TNC, PLAT, LAMC2, ACTG2, NR4A1, CYP2A6, KRT19, KRT18, COL5A1, DUOXA1, SCNN1A, FOSB, KCNN4, CD300LG, LTF, and MPZL2) out of 2316.

**Table 4:** Enriched terms obtained from a) ARCHS4 Tissues and b) NCI-60 Cancer Cell Lines via Enrichr according to produced genes by each method when using Dataset2. The best result is shown in bold.

| a) ARCHS4 Tissues |  |  |  |  |  |
| --- | --- | --- | --- | --- | --- |
| Method | Rank | Term | Overlap | P-value | Adjusted P-value |
| esvm | 21 | BREAST (BULK TISSUE) | **20/2316** | $1.00 \times 10^{-2}$ | $4.91 \times 10^{-2}$ |
| limma | 26 | BREAST (BULK TISSUE) | 11/2316 | $6.18 \times 10^{-1}$ | $9.99 \times 10^{-1}$ |
| sam | 11 | BREAST (BULK TISSUE) | 13/2316 | $3.72 \times 10^{-1}$ | $9.99 \times 10^{-1}$ |
| t-test | 17 | BREAST (BULK TISSUE) | 11/2316 | $6.18 \times 10^{-1}$ | $9.99 \times 10^{-1}$ |
| lasso | 6 | BREAST (BULK TISSUE) | 7/2316 | $1.03 \times 10^{-1}$ | $8.76 \times 10^{-1}$ |
| b) NCI-60 Cancer Cell Lines |  |  |  |  |  |
| Method | Rank | Term | Overlap | P-value | Adjusted P-value |
| esvm | - | MD-MB231 | - | - | - |
| | 4 | MCF7 | 6/397 | $1.47 \times 10^{-2}$ | $2.75 \times 10^{-1}$ |
| | 58 | HS578T | 1/176 | $5.87 \times 10^{-1}$ | $7.72 \times 10^{-1}$ |
| limma | - | MD-MB231 | - | - | - |
| | 43 | MCF7 | 2/397 | $5.93 \times 10^{-1}$ | $8.36 \times 10^{-1}$ |
|  | - | HS578T | - | - | - |
| sam | 32 | MD-MB231 | 2/150 | $1.72 \times 10^{-1}$ | $4.48 \times 10^{-1}$ |
| | 41 | MCF7 | 3/397 | $3.19 \times 10^{-1}$ | $6.46 \times 10^{-1}$ |
|  | - | HS578T | - | - | - |
| t-test | - | MD-MB231 | - | - | - |
| | 32 | MCF7 | 3/397 | $3.19 \times 10^{-1}$ | $6.80 \times 10^{-1}$ |
| | 49 | HS578T | 1/176 | $5.87 \times 10^{-1}$ | $8.43 \times 10^{-1}$ |
| lasso | 14 | MD-MB231 | 1/150 | $2.31 \times 10^{-1}$ | $6.54 \times 10^{-1}$ |
| | 33 | MCF7 | 1/397 | $5.04 \times 10^{-1}$ | $6.54 \times 10^{-1}$ |
| | 15 | HS578T | 1/176 | $2.66 \times 10^{-1}$ | $6.54 \times 10^{-1}$ |

The second-best method was sam with 13 expressed genes (DSP, IGFBP5, ODF3, TACSTD2, KLK8, SLC5A6, EFEMP1, FABP4, PER3, SLPI, CRTAC1, STAB1, and DACT2) out of 2316. Limma and *t*-test were tied for third-best performing methods with both having 11 expressed genes within the breast tissue.

Limma had the expressed genes, namely CAMSAP3, FABP4, JUP, TRAF4, SLFNL1, TRIM29, WNT9A, A2M, PCDH1, LOXL1, and SLC9A1 while *t*-test had these following genes: SLC22A23, MUCL1, GIPC3, PACS2, FZD7, PADI2, CLEC4F, CFB, KIAA0040, SYT7, and EGFR. The worst-performing method was lasso with 7 expressed genes (RECQL4, NR4A1, SLC44A4, GJD3, CLDN7, SHB, and LZTR1) out of 2316. Table 4(b) demonstrates enriched breast cancer cell lines terms from NCI-60 Cancer Cell Lines obtained via Enrichr. The best-performing method is esvm with a total of 7 expressed genes within MCF7 and HS578T breast cancer cell lines, distributed as follows. Six expressed genes (DCTN5, KRT19, DAAM1, KYNU, TRIM37, and MPZL2) out of 397 within MCF7 breast cancer cell line while 1 expressed gene (ACTG2) within the HS578T breast cancer cell line. In terms of MD-MB231 breast cancer cell line, esvm had no retrieved results. Therefore, results were designated as “-”. The second-best performing method was sam, resulted in 5 expressed genes within MD-MB231 and MCF7 breast cancer cell lines. Two expressed genes (RHEB and GARNL3) out of 150 were expressed within MD-MB231 breast cancer cell line, whereas 3 genes (PIAS3, DCTN5, and CTCF) out of 397 were expressed within MCF7 breast cancer cell line. For HS578T breast cancer cell line, no retrieved results were reported for sam (See “-”). The worst-performing method was lasso with 3 expressed genes within MD-MB231, MCF7, and HS578T breast cancer cell lines. These results demonstrate the good performance results of esvm in identifying breast tissues and cancer cell lines. In Supplementary DataSheet2_A, we include genes obtained from computational methods, provided to Enrichr to derive enrichment analysis results. Additionally, enrichment analysis results related to Table 4 are included in Supplementary Table2_A and Table2_B. Figure 4(a) displays the UpSet plot in terms of intersection lists of genes of all computational methods. Ninety-one unique genes were attributed to esvm. Sam, limma, *t*-test, and lasso had 94, 93, 92, and 33 unique genes, respectively. These results indicate that each method incorporated different computational steps resulted in different lists of genes. The number of intersected genes between each pair of computational methods is upper bounded by 4. In Supplementary DataSheet2_B, we include gene lists of computational methods related to Figure 4(a).

**Figure 4:**
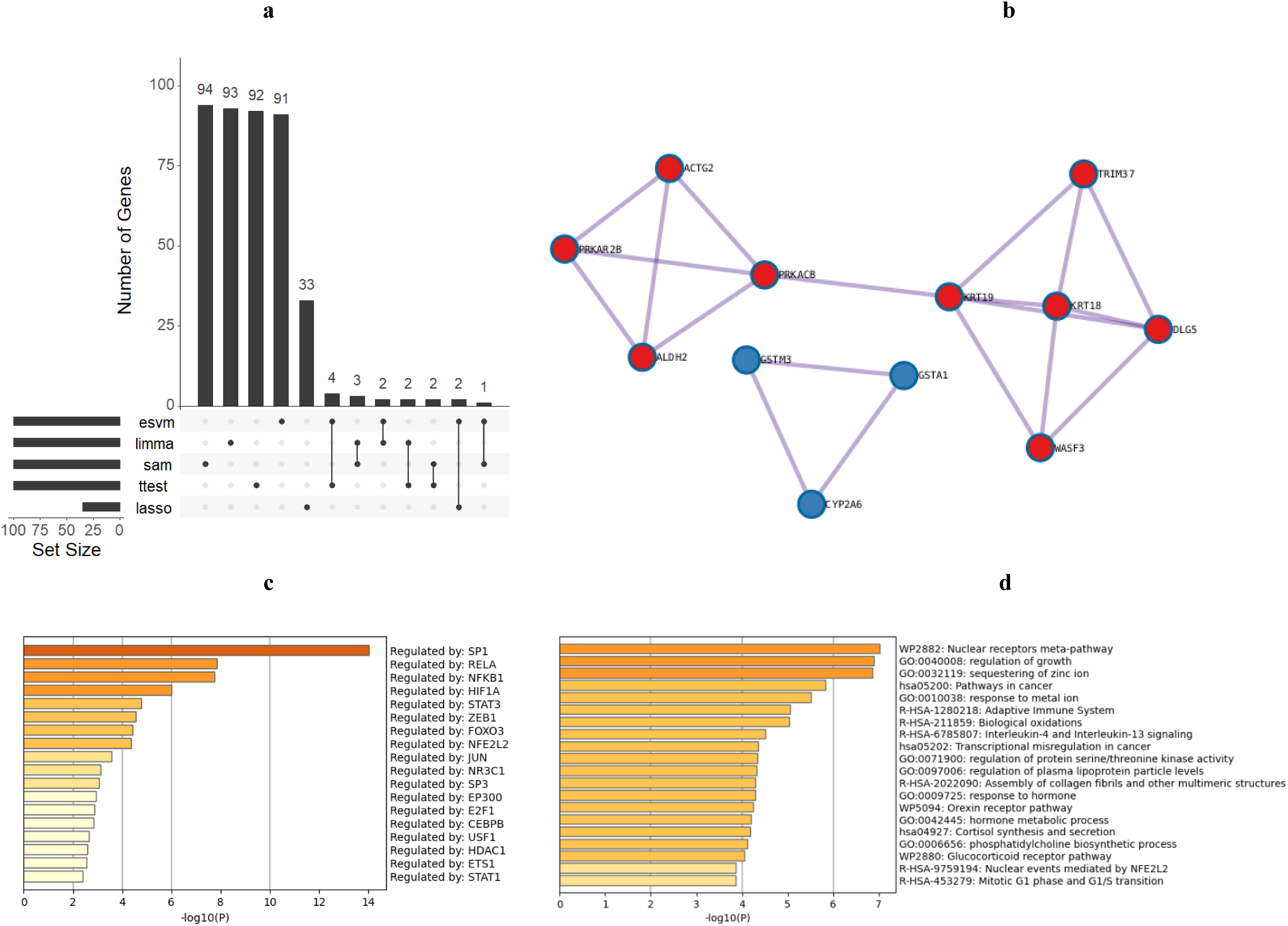
(a) UpSet plot of gene lists provided by the computational methods when using Dataset2. (b) Three clusters in a protein-protein interaction based on genes of esvm coupled with Metascape. (c) Eighteen transcription factors according to Metascape when coupled with genes from esvm. (d) Process and pathway enrichment analysis provided by Metascape according to the 100 genes of esvm.

As esvm performed better than baseline computational methods, we provided genes obtained from esvm to Metascape to unveil biological insights within breast cancer drug response. Figure 4(b) reports the following 12 genes obtained from PPI network: ACTG2, PRKAR2B, ALDH2, PRKACB, KRT19, KRT18, WASF3, DLG5, TRIM37, GSTM3, GSTA1, and CYP2A6. Four genes (KRT18, KRT19, PLCB4, and PRKACB) were related to Estrogen signaling pathway, which is reported to play a key role in breast cancer progression and treatment [67–69]. Three genes (CYP2A6, GSTA1, and GSTM3) were linked to chemical carcinogenesis - DNA adducts pathway, involved in cancer development [70, 71]. In Figure 4(c), we report the following 18 transcription factors: SP1, RELA, NFKB1, HIF1A, STAT3, ZEB1, FOXO3, NFE2L2, JUN, NRC1, SP3, EP300, E2F1, CEBPB, USF1, HDAC1, ETS1, and STAT1.

STAT3 is involved in breast cancer progression [72]. E2F1 and EP300 have been reported to be involved in breast cancer development and metastasis [73–75]. These results demonstrate the importance of these TFs and such mutations or alterations can affect gene regulation and thereby contribute to breast cancer development. In Figure 4(d), the top enriched term was nuclear receptors meta-pathway. Ten genes (CYP2A6, GSTA1, GSTM3, HMOX1, ME1, S100P, SCNN1A, ABCC4, PLK2, and B3GNT5) were linked to nuclear receptors meta-pathway, which has been related to breast cancer cell growth via nuclear receptors such as estrogen receptors [76]. Moreover, the 13 genes (AGT, BCL6, CDKN2C, FGFR3, GJA1, TNC, MT1X, S100A8, S100A9, SYT1, SOCS2, SEMA3C, and CHPT1) were related to regulation of growth process, which is linked to breast cancer cell growth. Eleven genes (AGT, BIRC5, CCND2, FGFR3, GSTA1, GSTM3, HMOX1, CXCL8, LAMC2, PLCB4, and PRKACB) were related to Pathways in cancer, which is linked to breast cancer development and metastasis [77]. Six genes (BCL6, CCND2, CDKN2C, CXCL8, PLAT, and PROM1) were related to transcriptional misregulation in cancer, which is linked to mutations and altered gene expression in breast cancer [78]. Enrichment analysis results obtained from Metascape are provided in Supplementary Enrichment_Dataset2.

Table 5 shows drugs and drug targets within IDG Drug Targets 2022. Hydroxycarbamide and gemcitabine are ribonucleotide reductase enzyme (RNR) inhibitors that have been used for breast cancer treatment (See Figure 5) [79, 80]. The two drugs are associated with RRM2 as a drug target [81]. Daunorubicin inhibits DNA replication and Cyclophosphamide causing damage to the DNA of cancer cells and thereby causing cancer cells to die. TOP2A was a drug target for Daunorubicin, whereas RRM2 was a drug target for Cyclophosphamide. In Supplementary Table2_C, we report enrichment analysis results for IDG Drug Targets 2022.

**Table 5:**
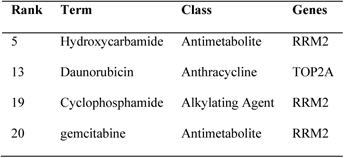
Retrieved enriched terms from IDG Drug Targets 2022 via Enrichr according to uploaded genes from esvm.

**Figure 5:**
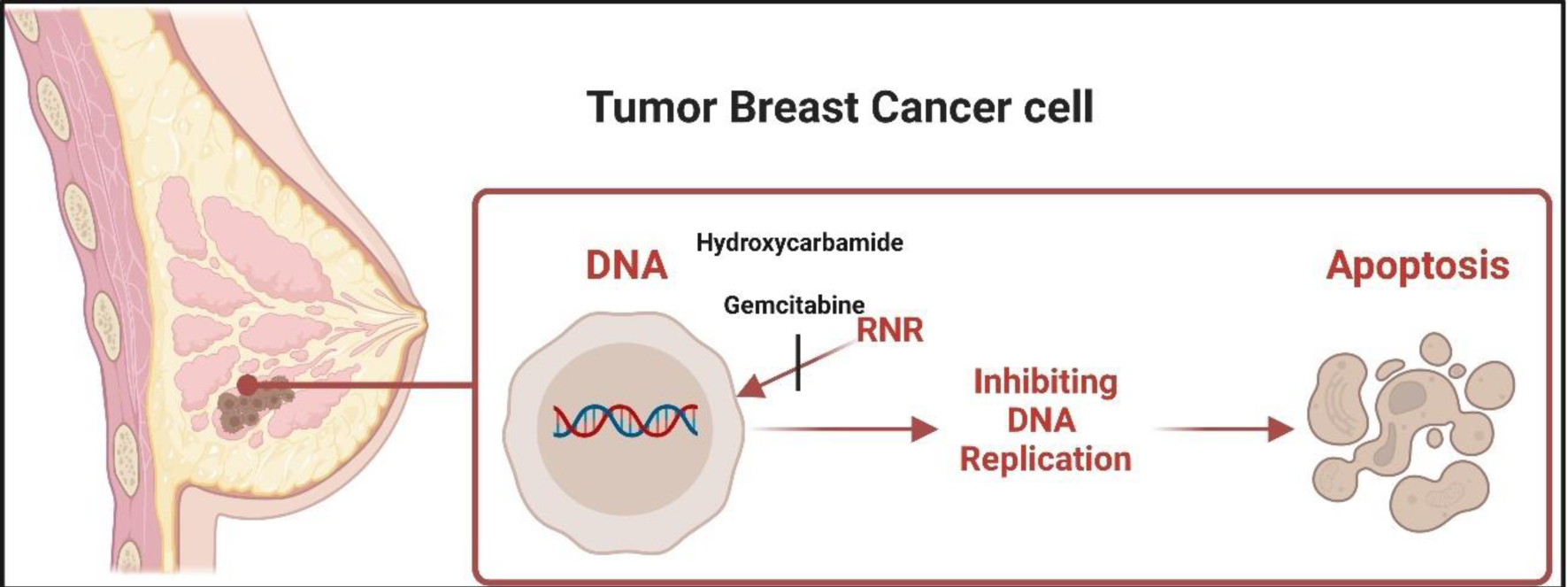
Hydroxycarbamide and gemcitabine inhibiting ribonucleotide reductase (RNR) enzyme, which inhibits the DNA replication during the cell cycle and inducing apoptosis.

#### 3.2.3 Dataset3

After uploading produced genes via each method to Enrichr, we show retrieved breast tissue terms within ARCHS4 Tissues (See Table 6(a)) and breast cancer cell lines terms within NCI-60 Cancer Cell Lines (See Table 6(b)). From Table 6(a), we can see that esvm generated the best results. Particularly, esvm had 6 expressed genes (CSF1, CCND1, ERBB3, IRS1, ERBB2, and ESR1) out of 2316 within breast tissue, followed by sam and *t*-test, both having 4 common expressed genes (RET, CCND1, IRS1, and ERBB2) out of 2316 within breast tissue. Lasso had 2 (ESR1 and EGFR) expressed genes out of 2316 within breast tissue while limma was the worst-performing method having 1 expressed gene (ABL1) out of 2316 within breast tissue. In Table 6(b), esvm is also the best-performing method by having 3 expressed genes out of 397 within MCF7 breast cancer cell line. No results were associated with MD-MB231 and HS578T breast cancer cell lines. Therefore, we indicated results by “-“. Both sam and *t*-test were tied by having 2 expressed genes out of 397 within MCF7 breast cancer cell line. As esvm, no reported results were found for the other two breast cancer cell lines (i.e., MD-MB231 and HS578T). The worst-performing method was limma where no reported results were found for the three breast cancer cell lines. These results demonstrate the superiority of esvm when identifying breast cancer tissue and cell lines. We include enrichment analysis results regarding Table 6 in Supplementary Table3_A and Table3_B.

**Table 6:** Enriched terms obtained from a) ARCHS4 Tissues and b) NCI-60 Cancer Cell Lines via Enrichr according to produced genes by each method when using Dataset3. The best result is shown in bold.

| a) ARCHS4 Tissues |  |  |  |  |  |
| --- | --- | --- | --- | --- | --- |
| Method | Rank | Term | Overlap | P-value | Adjusted P-value |
| esvm | 3 | BREAST (BULK TISSUE) | **6/2316** | $5.28 \times 10^{-1}$ | $9.97 \times 10^{-1}$ |
| limma | 40 | BREAST (BULK TISSUE) | 1/2316 | $9.78 \times 10^{-1}$ | $9.78 \times 10^{-1}$ |
| sam | 4 | BREAST (BULK TISSUE) | 4/2316 | $2.99 \times 10^{-1}$ | $9.47 \times 10^{-1}$ |
| t-test | 4 | BREAST (BULK TISSUE) | 4/2316 | $2.99 \times 10^{-1}$ | $9.47 \times 10^{-1}$ |
| lasso | 9 | BREAST (BULK TISSUE) | 2/2316 | $4.12 \times 10^{-1}$ | $7.71 \times 10^{-1}$ |
| b) NCI-60 Cancer Cell Lines |  |  |  |  |  |
| Method | Rank | Term | Overlap | P-value | Adjusted P value |
| esvm | - | MD-MB231 | - | - | - |
| | 3 | MCF7 | 3/397 | $7.68 \times 10^{-2}$ | $5.88 \times 10^{-1}$ |
|  | - | HS578T | - | - | - |
| limma | - | MD-MB231 | - | - | - |
|  | - | MCF7 | - | - | - |
|  | - | HS578T | - | - | - |
| sam | - | MD-MB231 | - | - | - |
| | 3 | MCF7 | 2/397 | $8.14 \times 10^{-2}$ | $4.41 \times 10^{-1}$ |
|  | - | HS578T | - | - | - |
| t-test | - | MD-MB231 | - | - | - |
| | 3 | MCF7 | 2/397 | $8.14 \times 10^{-2}$ | $4.41 \times 10^{-1}$ |
|  | - | HS578T | - | - | - |
| lasso | - | MD-MB231 | - | - | - |
| | 18 | MCF7 | 1/397 | $2.13 \times 10^{-1}$ | $3.08 \times 10^{-1}$ |
|  | - | HS578T | - | - | - |

In Figure 6(a), we show the UpSet plot showing intersection of produced genes among all methods when Dataset3 is used. From the leftmost, it appears that esvm differs from all other methods by having unique 20 unique genes. Limma and lasso have 17 and 6 unique genes, respectively. Esvm, sam and *t*-test share 13 genes. Limma and esvm share 12 genes. Sam and *t*-test share 8 genes. Lasso and esvm share 3 genes. Sam, *t*-test, and lasso share 2 genes. It can also be seen that the number of common genes between the remaining intersection of methods is 1. These results demonstrate that our method is different from remaining methods, attributed to different computational steps involved in the computations of esvm. In Supplementary DataSheet3_B, we report genes related to UpSet plot. Figure 6(b) reports the following 19 genes obtained from PPI network: AR, BIRC5, CCND1, RB1, STAT1, STAT3, ESR1, IRS1, PTEN, ERBB2, ERBB3, ALK, AKT1, MET, JAK2, IGFIR, EGFR, TP53, and MTOR. Thirteen genes (AKT1, ARAF, CCND1, EGFR, ERBB2, ERBB3, MTOR, IGF1R, JAK2, MET, PTEN, RAF1, and STAT3) were linked to EGFR tyrosine kinase inhibitor resistance, involved in resistance mechanism of EGFR inhibitor, and thereby breast cancer progression [82]. Twelve genes (AKT1, CCND1, ERBB2, ESR1, MTOR, IRS1, JAK2, NOS3, PTEN, RAF1, STAT1, and STAT3) were involved in leptin signaling pathway, which is related to breast cancer malignancy [83, 84]. Five genes (BIRC5, CCND1, ESR1, RB1, and CHEK2) were involved in PID FOXM1 PATHWAY, which has been reported as a crucial oncogenic transcription factor that promotes breast cancer progression and growth [85, 86].

**Figure 6:**
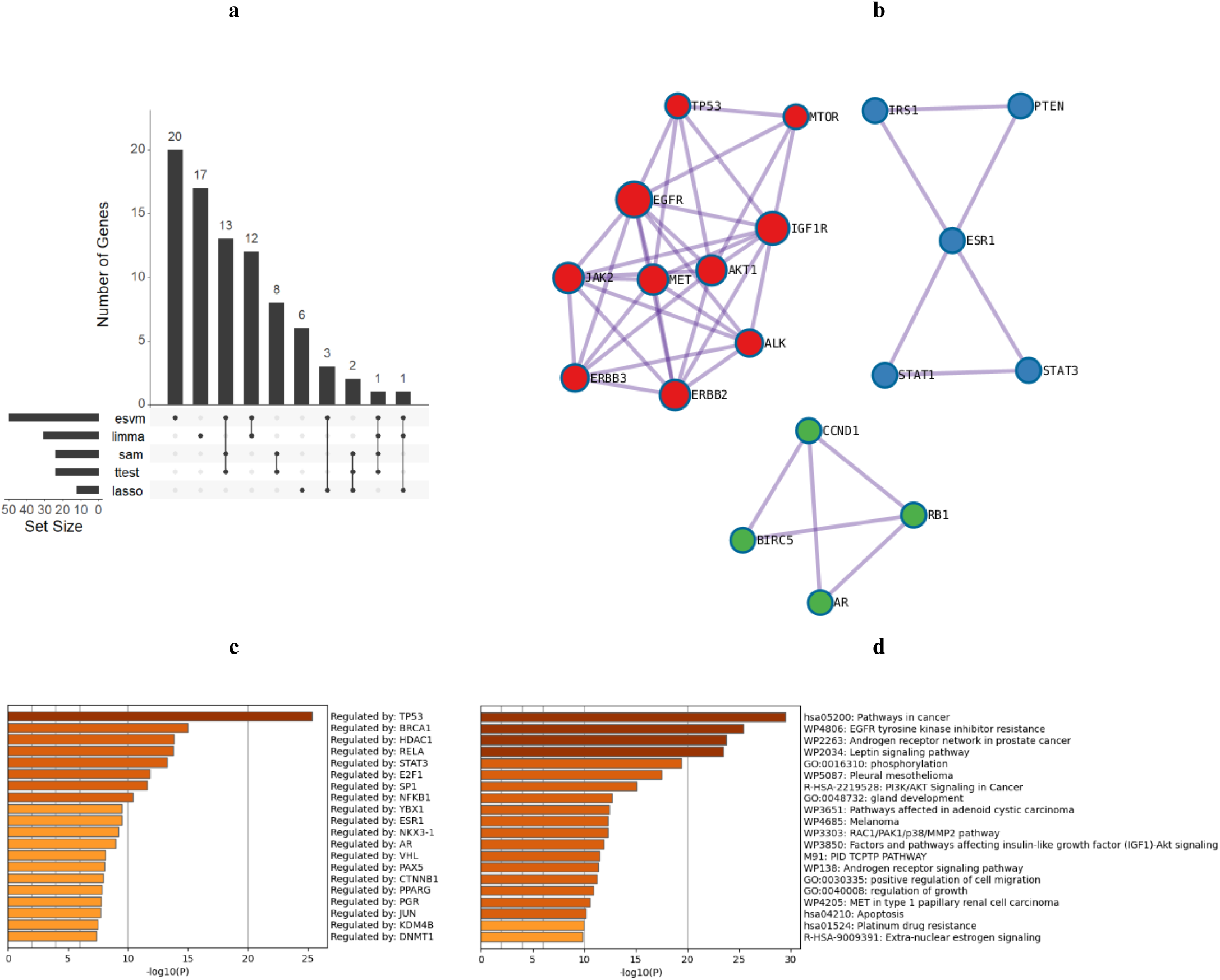
(a) UpSet plot of gene lists provided by the computational methods when using Dataset3. (b) Three clusters in a protein-protein interaction based on genes of esvm coupled with Metascape. (c) Twenty transcription factors according to Metascape when coupled with genes from esvm. (d) Process and pathway enrichment analysis provided by Metascape according to the genes of esvm.

In Figure 6(c), we report the following 20 transcription factors (TFs): TP53, BRCA1, HDAC1, RELA, STAT3, E2F1, SP1, NFKB1, YBX1, ESR1, NKX3-1, AR, VHL, PAX5, CTNNB1, PPARG, PGR, JUN, KDM4B, and DNMT1. Interestingly, TP53 mutation has been reported to be the most frequently occurring in breast cancer [87, 88]. Patients with BRCA1 mutations are at higher risk of developing breast cancer and thereby considered as an important biomarker [89, 90]. HDAC1 has been reported to be related to breast cancer cell proliferation [91]. These results demonstrate the importance of these TFs and can aim in developing therapeutic strategies for breast cancer treatment. Figure 6(d) reports top enriched terms regarding processes and pathways. Twenty-one genes (AKT1, ALK, BIRC5, AR, ARAF, CCND1, CASP7, EGFR, ERBB2, ESR1, MTOR, IGF1R, JAK2, MET, PTEN, RAF1, RB1, STAT1, STAT3, TP53, and FADD) were linked to Pathways in cancer (hsa05200), coinciding with recently reported results as one of the top enriched pathways in breast cancer [92].

Thirteen genes (AKT1, ARAF, CCND1, EGFR, ERBB2, ERBB3, MTOR, IGF1R, JAK2, MET, PTEN, RAF1, and STAT3) were related to EGFR tyrosine kinase inhibitor resistance (WP4806), which was reported as significant pathway associated with breast cancer [78, 93]. Twelve genes (AKT1, CCND1, ERBB2, ESR1, MTOR, IRS1, JAK2, NOS3, PTEN, RAF1, STAT1, and STAT3) were linked to Leptin signaling pathway (WP2034), reported to have a key role in breast cancer tumorigenesis [94]. Nine genes (AKT1, EGFR, ERBB2, ERBB3, ESR1, MTOR, IRS1, MET, and PTEN) were linked to PI3K/AKT Signaling in Cancer (R-HSA-2219528) in which its inactivation suppressing the proliferation of breast cancer cells and thereby inducing apoptosis [95]. Seven genes (AKT1, BIRC5, ATM, CASP7, RAF1, TP53, and FADD) were related to apoptosis (hsa04210), which plays a key role in controlling the excessive proliferation of breast cancer cells [96]. In Supplementary Enrichment_Dataset3, we include enrichment analysis results obtained from Metascape pertaining to Dataset3.

Table 7 reports drug terms and drug targets within IDG Drug Targets 2022. Ceritinib aims to inhibit anaplastic lymphoma kinase (ALK) enzyme and thereby blocking the ability of tumors to grow and promoting apoptosis (See Figure 7) [97]. Erlotinib inhibits the effect of tyrosine kinase enzyme on epidermal growth factor receptor (EGFR) and thereby preventing proliferation of cancer cells and inducing apoptosis. In Supplementary Table3_C, we report enrichment analysis results regarding drug terms within IDG Drug Targets 2022.

**Table 7:**
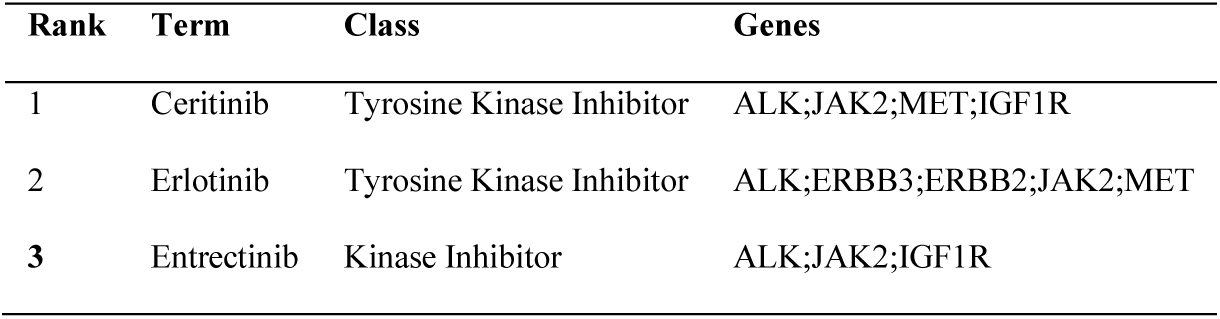
Retrieved enriched terms from IDG Drug Targets 2022 via Enrichr according to uploaded genes using Dataset3, showing genes (column: Genes) associated with drugs (column: Term). Rank column shows the order of terms when retrieved.

| Rank | Term | Class | Genes |
| --- | --- | --- | --- |
| 1 | Ceritinib | Tyrosine Kinase Inhibitor | ALK;JAK2;MET;IGF1R |
| 2 | Erlotinib | Tyrosine Kinase Inhibitor | ALK;ERBB3;ERBB2;JAK2;MET |
| 3 | Entrectinib | Kinase Inhibitor | ALK;JAK2;IGF1R |

**Figure 7:**
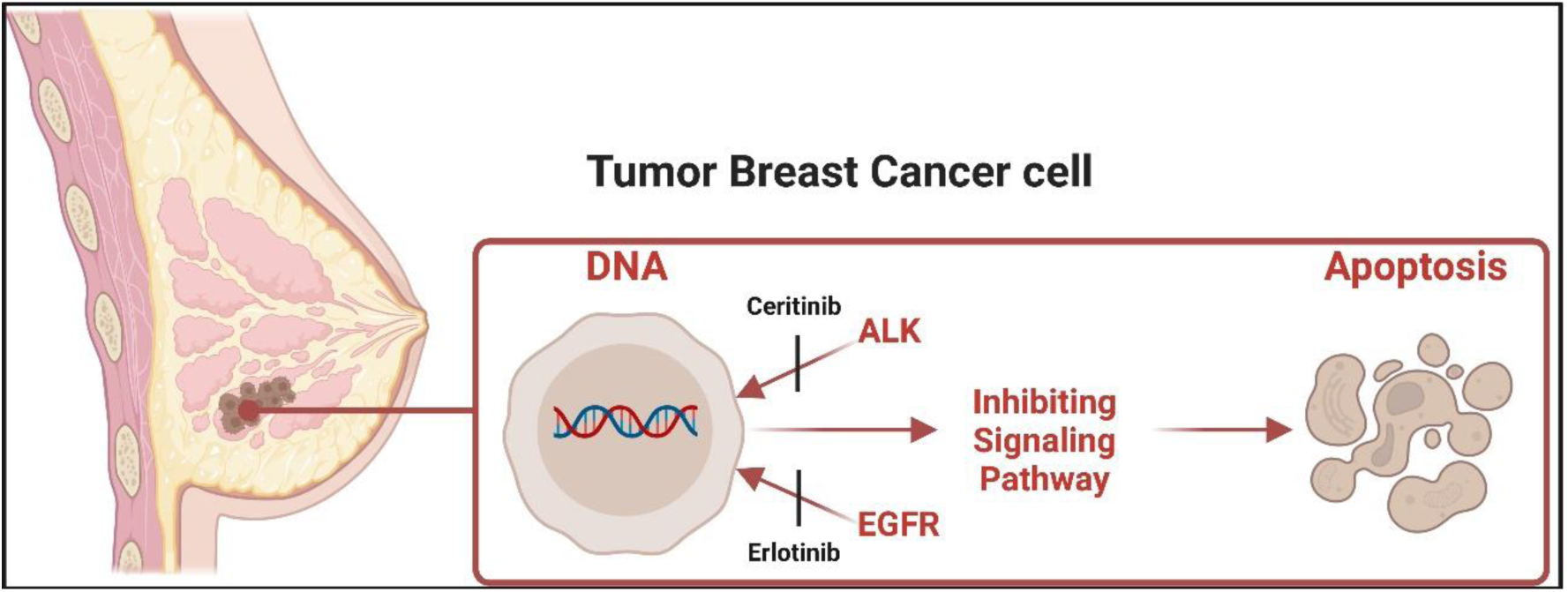
Ceritinib and erlotinib are anaplastic lymphoma kinase (ALK) and epidermal growth factor receptor (EGFR) inhibitors, receptively.

### 3.3 Models Introspection

## Dataset1

In Figure 8, we aim to get computational insights pertaining to studied learning-based models applied in the Results section. Figure 8(a) demonstrated that our method esvm leads to non-zero weights while lasso in Figure 8(b) leading to a sparser representation and thereby many zero coefficients, attributed to the L1 penalty as in [28]. In Figure 8(c), it can be seen that lasso is ∼ 6.14 × faster than esvm. Figures 8(d and e for esvm and lasso, respectively) demonstrate that prediction differences between breast cancer patients achieving pathological complete response against those having residual disease were statistically significant (*p-*value of all models <2.2 × 10^−16^, obtained from a *t*-test).

**Figure 8:**
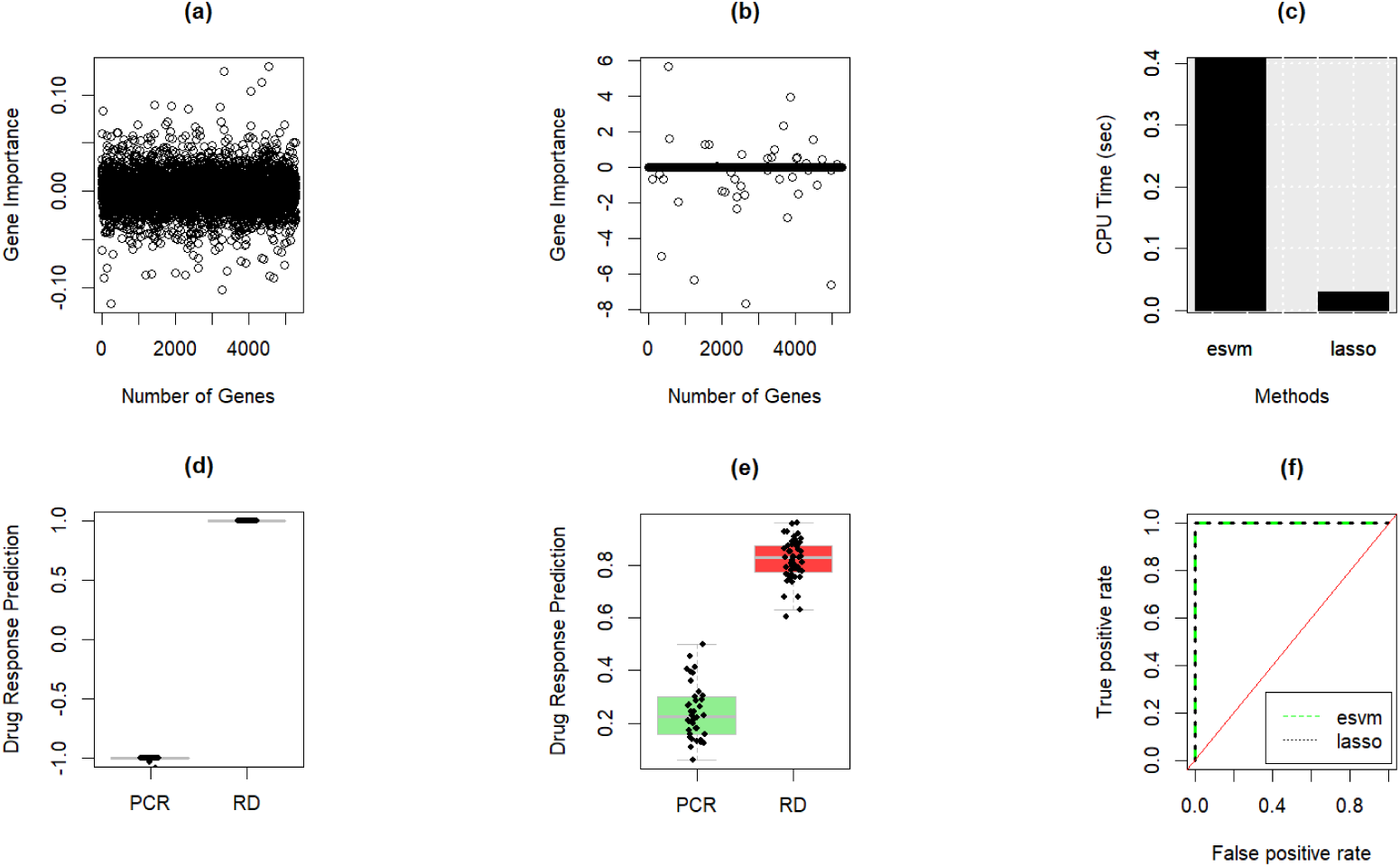
Classification models comparison for predicting drug response of TCH, TCHTy, and TCTy in breast cancer (BC) patients when Dataset1 is used. Gene importance when esvm (a) and lasso (b) are applied. Computational running time (c) for esvm and lasso. Boxplot and strip chart of drug sensitivity prediction for BC patients using esvm (d) and lasso (e). ROC curve (f) demonstrating the prediction performance. TCH is docetaxel, carboplatin, and trastuzumab. TCHTy is docetaxel, carboplatin, trastuzumab and lapatinib. TCTy is docetaxel, carboplatin and lapatinib. PCR is pathological complete response. RD is residual disease.

These results show that both induced models using Dataset1 are expected to be general predictors for drugs with PCR and RD responses. For the ROC curves of esvm and lasso in Figure 8(f), both models achieved an area under the ROC curve (AUC) of 1.00.

## Dataset2

As shown in Figure 9, different aspects related to models are reported as follows. Figure 9(a) shows that esvm leading to non-sparse representation in which the weight vector w consisting of non-zero weights. On the other hand, Figure 9(b) demonstrates that lasso leading to a sparse representation in which many of the coefficients β are zeros. Figure 9(c) displays that lasso is 44 × faster than esvm.

**Figure 9:**
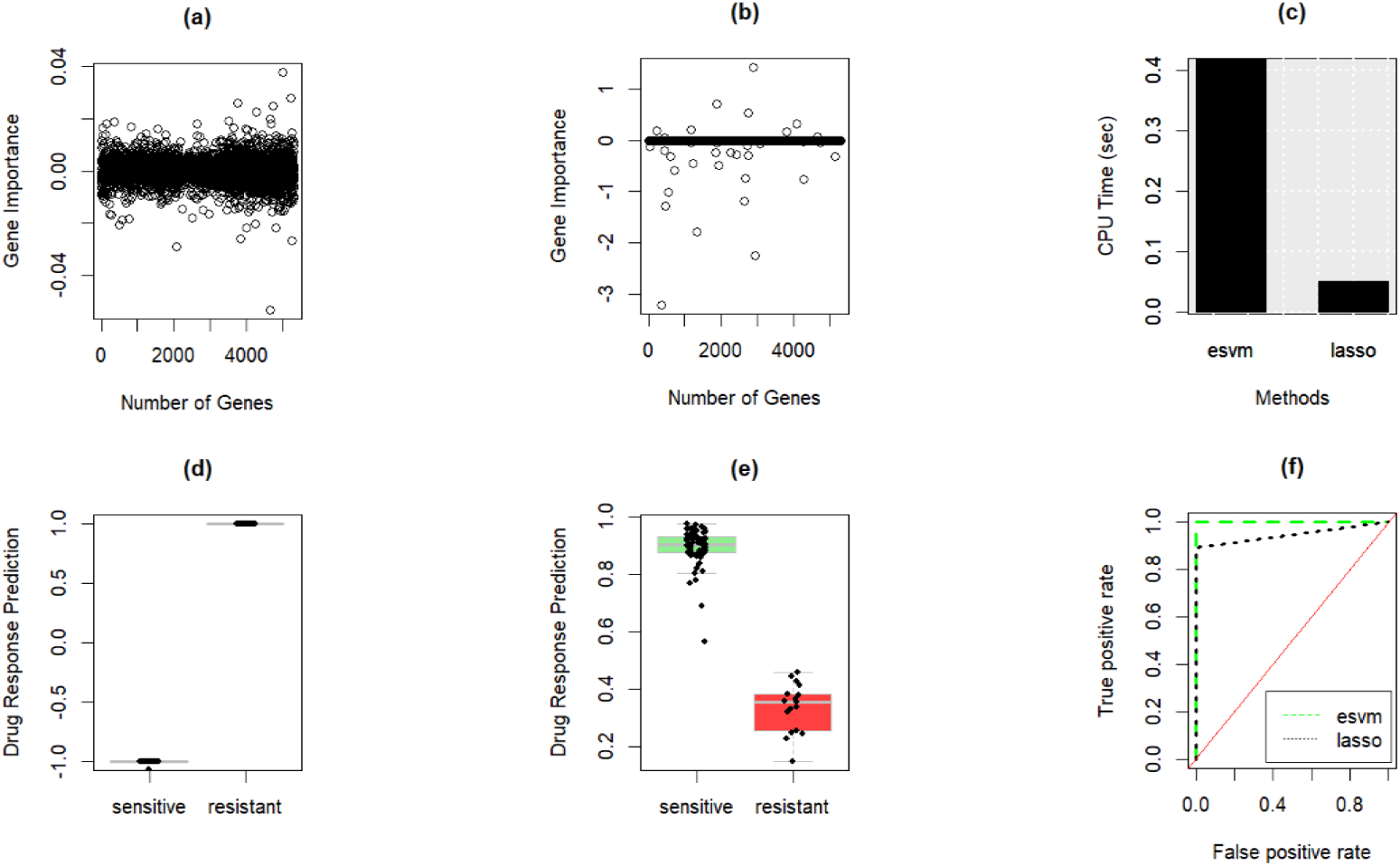
Classification models comparison for predicting TFEC drug response of in breast cancer (BC) patients when Dataset2 is used. Gene importance when esvm (a) and lasso (b) are applied. Computational running time (c) for esvm and lasso. Boxplot and strip chart of drug sensitivity prediction for BC patients sensitive against those resistant to the drug treatment using esvm (d) and lasso (e). ROC curve (f) demonstrating the prediction performance. TFEC is docetaxel, 5-fluorouracil, epirubicin, and cyclophosphamide.

Figure 9 (d) for esvm demonstrates that prediction differences between sensitive breast cancer patients against those having resistant drug response were statistically significant (*p-*value < 2.2 × 10^−16^ , obtained from a *t*-test). For lasso in Figure 9 (e), prediction differences between the two groups (i.e., BC patients sensitive to a drug against those having resistant response) were not statistically significant (*p-* value =1, obtained from a *t*-test). These obtained results show that esvm is expected to be a general predictor for drug sensitivity and resistance. Lasso tends to be a specific predictor. Figure 9(f) displays the ROC curves for esvm and lasso, where the former has an AUC of 1.00 and the latter has an AUC of 0.947.

## Dataset3

Figure 10 reports various computational aspects for studied models from classification perspective. It can be easily noticed that esvm doesn’t shrink weights w to zero (Figure 10(a)) while lasso has sparser representation attributed to the L1 regularization shrinking coefficients β to zero (See Figure 10(b)). In terms of efficiency, Figure 10(c) shows that that lasso is 1974 × faster than esvm. In terms of generalization, Figure 10(d and e) demonstrates that prediction differences of esvm and lasso between breast cancer patients achieving complete response (CR) against those of failed complete response (FCR) were statistically significant (*p-*value <2.2 × 10^−16^, obtained from a *t*-test). However, esvm had a better AUC of 0.722 while lasso had a lower AUC of 0.555 (See Figure 10(f)).

**Figure 10:**
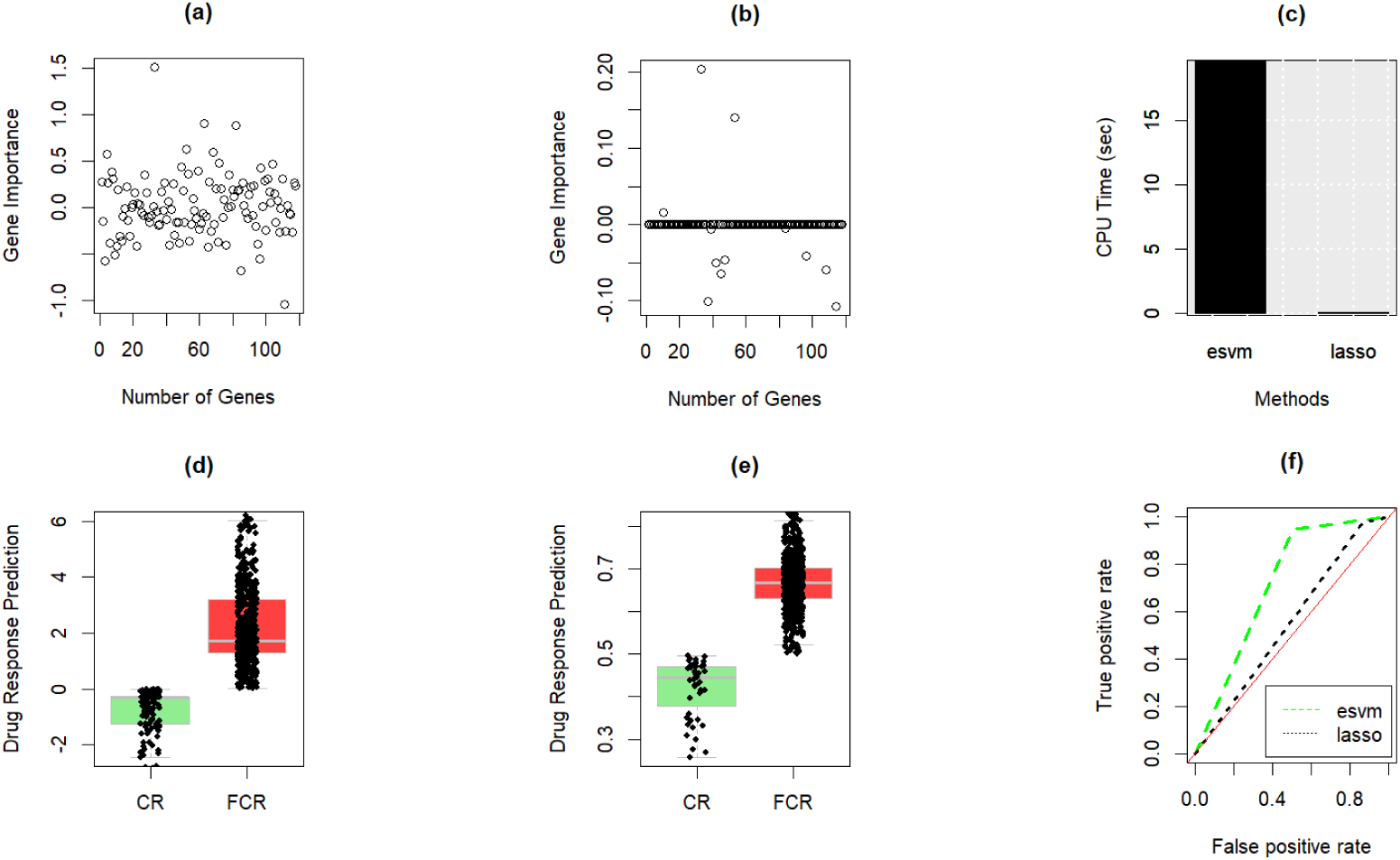
Classification models comparison for predicting multiple drug combinations response in breast cancer (BC) patients when Dataset3 is used. Gene importance when esvm (a) and lasso (b) are applied. Computational running time (c) for esvm and lasso. Boxplot and strip chart of drug sensitivity prediction for BC patients sensitive against those resistant to the drug treatment using esvm (d) and lasso (e). ROC curve (f) demonstrating the prediction performance.

These performance results indicate that prediction performance difference between esvm and lasso is 16.7% when AUC is considered.

## Scalability

In Figure 11(a-j), we report the computational running time for increased dimensionality starting from 50000 dimensions to 500000 dimensions and fixing the number of rows to 100. The generation of X was done according to the uniform distribution 𝑈(0,1), Y was generated in which we balanced the label distribution. When the number of dimensions is 50000 (See Figure 11(a)), lasso was 27 and 121.5 × faster than esvm and svm, respectively. Our method esvm was 4.5 faster than the baseline svm. The average running times for esvm, svm, and lasso spanning over all results in Figure 11(a-j) are 3.54 seconds, 23.88, and 0.275, respectively. That means our method, esvm, on average is 6.74 × faster than svm while lasso was 12.87 and 86.83 × faster than esvm and svm, receptively. These results demonstrate the computational efficiency of our method over the svm implementation using e1071 package in R. In Supplementary Running_Time, we include all running time results related to Figure 11(a-j).

**Figure 11:**
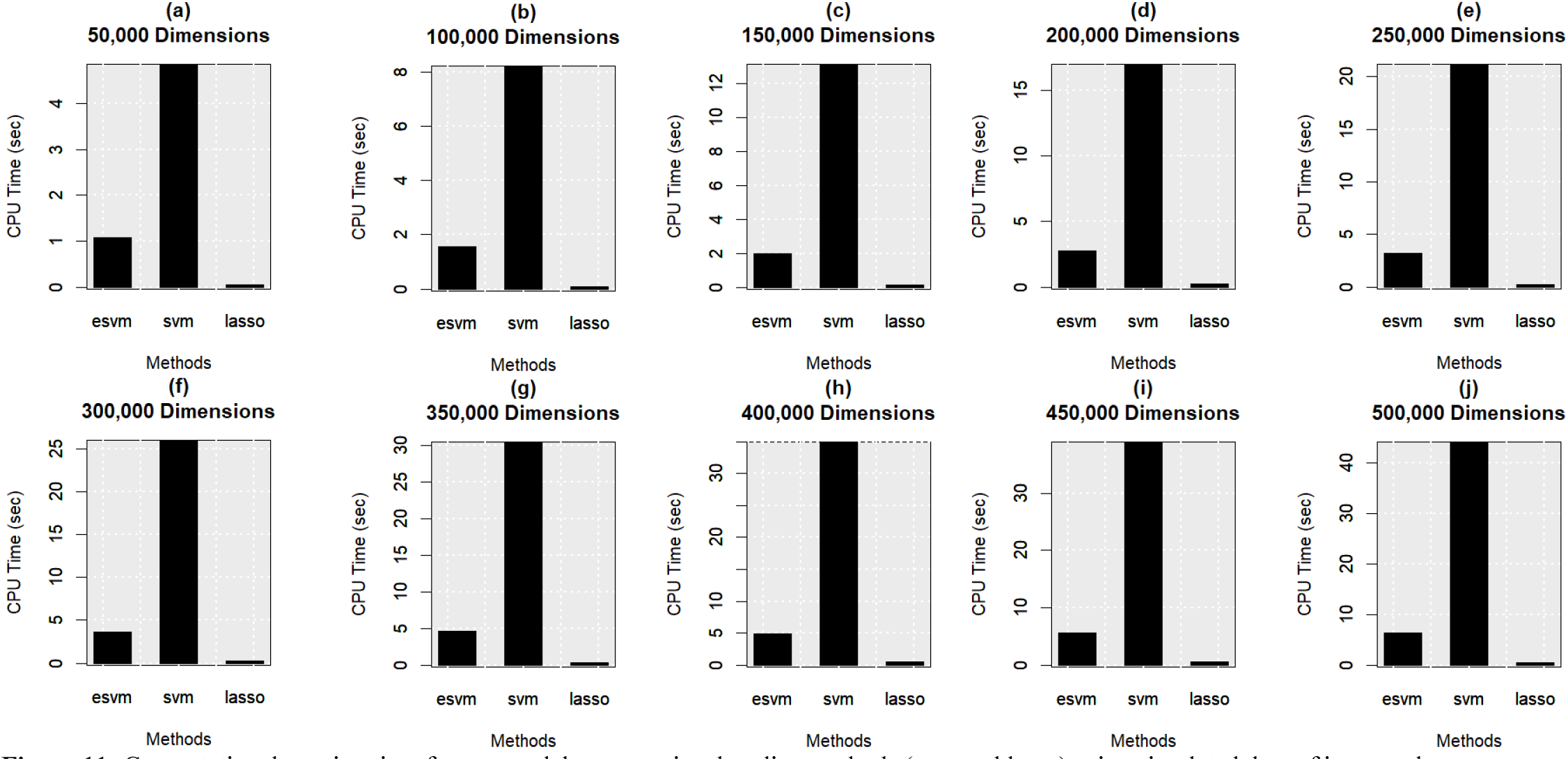
Computational running time for our model esvm against baseline methods (svm and lasso) using simulated data of increased dimensionality.

## 4 Discussion

Identifying critical genes, drugs, drug targets, and transcription factors play a key role in unveiling the underlying drug response mechanism of breast cancer. Therefore, we introduced an AI-based computational framework functioning as follows. First, because gene expression datasets consist of many genes compared to the number of samples, the optimization problem formulation of svm in terms of finding weight vector w and bias term *b* is impractical (See Equation 1). Therefore, we employ the dual form of SVM optimization problem formulated in terms of finding lambda λ as shown in Equation 2. Then, we recover the weight vector w using Equation 3. Our method, esvm, takes as input gene expression data along with breast cancer drug responses, obtained from the GEO database. The output is a list of selected 100 genes according to the top 100 corresponding weights in w. Then, we performed enrichment analysis via providing the output genes to two enrichment analysis tools, Enrichr and Metascape. When Enrichr is considered, our method outperformed the baseline methods via having more expressed gene in breast cancer cell lines, including MD-MB231, MCF7, and HS578T. Moreover, our method esvm had more expressed genes in breast tissue. For Metascape, our method identified important genes including tumor suppressor genes (e.g., TP53 and BRCA1), TFs, drugs, drug targets, pathways and biological processes that play a key role in understanding breast cancer drug response mechanism.

As computational running time plays a key role in the gene selection process, our method esvm was way faster than the baseline svm. Although lasso was the fastest method, esvm had more expressed genes in breast cancer cell lines as well as breast tissues when compared to lasso. Therefore, the improvements in esvm are attributed to (1) superiority of computational efficiency when compared to the baseline svm; and (2) high performance results measured using the AUC when compared to lasso. Another advantage of esvm attributed to the computational efficiency is the computational feasibility to explicitly change the data representation and applying esvm to identify important genes. As a results, esvm can aid in analyzing gene expression data coupled with clinical data, and other profiling datasets.

Drug responses in this study were classified as follows. For Dataset1, the treated groups were classified during phase II trial as (1) pathological complete response referring to the absence of invasive cancer in axilla and breast; and (2) residual disease referring to the presence of invasive cancer in breast and axilla. In terms of Dataset2, treated groups after neoadjuvant therapy within Phase II multicenter trial were classified as (1) sensitive referring to patients completely responding to the treatment; and (2) resistant referring to not completely responding to the treatment. Regarding Dataset3, the treated groups were categorized during neoadjuvant I-SPY2 trial as (1) complete response when responding to the treatment; and (2) failed complete response when not completely responding to the treatment.

We specify parameters of methods in our study for gene selection as follows. esvm(*C* = 2), lasso(λ=0.05), and svm (setting parameters associated with the linear kernel to their default values). Each method producing a gene list in which the probability of each method coinciding with our method esvm is estimated as follows. For Dataset1, Dataset2 and Dataset3, the probabilities are equal to , respectively; (^𝑛^) indicates the binomial coefficient in which *n* is the total number of genes in the considered dataset and *k* is to the total number of genes produced by esvm. Therefore, the odds of having a method producing results like ours are unlikely to occur.

We evaluated the performance of selected genes from a biological perspective against deep learning methods, including DeepLIFT, DeepSHAP, and LRP. In the three datasets (see Supplementary Additional file: Tables S1(b), S2(b), and S3(b)), our method had more expressed genes in breast cancer cell lines (MD-MB231, MCF7, and HS578T). Specifically, in Dataset1, esvm had a total of eight expressed genes compared to six (two and four) for DeepLIFT (DeepSHAP and LRP). For Dataset2, esvm had a total of seven expressed genes while DeepLIFT, DeepSHAP, and LRP had a total of five, five, and six, respectively. In terms of Dataset3, esvm had a total of three expressed genes while each deep learning methods had a total of two expressed genes. In our study, we identified genes enriched in terms related to breast cancer cell lines and our method was the best. On the other hand, when considering genes in breast tissue irrespective to cell types, DeepLIFT was the best (see Supplementary Additional file: Tables S1(a), S2(a), and S2(a)). As a fraction of genes is expressed in each breast cancer cell line, our method had more expressed genes related to studied breast cancer cell lines. Therefore, these results demonstrate the superiority of esvm. Additional details for studied deep learning models are in Supplementary Additional file (Figures S1 and S2) and Supplementary DLModels.

To further demonstrate the effectiveness of esvm as a gene selection method, we assessed the performance from a classification perspective of our method against adapted deep learning methods for gene selection including Deep Learning Important FeaTures (DeepLIFT) [8], Deep SHapley Additive exPlanations (DeepSHAP) [9], and Layer-wise relevance propagation (LRP) [10]. Tables S4-S6 in Supplementary Additional file demonstrate the performance results (and standard deviation) when SVM is coupled with each gene set produced by each method for the three datasets. It can be seen from Table S4 in Supplementary Additional file that when SVM is coupled with Dataset1 of selected genes via esvm generated the highest accuracy (ACC) of 0.921, the highest balanced accuracy (BAC) of 0.935, the highest Matthews correlation coefficient (MCC) of 0.849. For Dataset2 (see Table S5 in Supplementary Additional file), SVM when coupled with gene set via esvm in Dataset2 generated the highest ACC of 0.978, the highest BAC of 0.960, the highest F1 of 0.950, and highest MCC of 0.944. The same holds true for Dataset3 in which SVM when coupled with dataset of genes selected from esvm achieved the highest performance results based on the four performance measures (see Table S6 in Supplementary Additional file).

When evaluating the performance from a classification perspective for the three datasets using lasso (rather than SVM) as a learning algorithm (see Supplementary Additional file: Tables S7-S9), Table S7 in Supplementary Additional file demonstrates that lasso when coupled with Dataset1 of genes selected via esvm generated the highest ACC of 0.662, the highest BAC of 0.660, the highest F1 of 0.738. In terms of Dataset2 (see Table S8 in Supplementary Additional file), lasso when coupled with Dataset2 with selected genes from esvm generated the highest BAC of 0.537 while other methods had marginal improvements over our method (esvm) when ACC performance measure was considered. The same holds true for Dataset3 in which lasso when coupled with Dataset3 of genes from esvm achieved the highest performance results using two performance metrics in imbalanced classification. Particularly, lasso with genes from esvm achieving the highest BAC of 0.627, and the highest MCC of 0.280. These results demonstrate the effectiveness of our method in exploring discriminatory genes. Combined confusion matrices for performance results using SVM and lasso are provided in Supplementary Additional file, displayed in Figures S3-S8 and the sum of entries in each confusion matrix is equal to the number of samples in the corresponding dataset.

In terms of the number of common genes in Dataset1, it can be seen from Figure 2(a) that the number of common genes (if exists) between any pair of methods is not exceeding 2. For example, limma and sam had two common genes (ATP2B1 and RNF186), esvm and *t-*test had two common genes (RAB38 and ABHD1), and sam and *t-*test had two common genes (MFN2 and RPS24). The rest of intersections are provided in Supplementary DataSheet1_B. For Dataset2, Figure 4(a) shows that esvm and *t-*test had four common genes (PTCHD1, CTSB, ALDH2, and CHGB), limma and sam had three common genes (FABP4, SNX32, and GBP6). In Supplementary DataSheet2_B, we include the rest of common genes. For Dataset3, Figure 6(a) shows that esvm, sam, and *t-*test had 13 common genes (CCND1, EGFR.3, EIF4GI,ERBB2, ERBB3.1, ESR1.1, IGF1R, IRS1, JAK2, KS6B1.2, PD1L1.3, STAT1, and STA5A).

The rest of intersections are provided in Supplementary DataSheet3_B. In Dataset1, Figures. 2(b-d) are presented in tabular format in Tables S10-S12 in Supplementary Additional file. For Dataset2, the corresponding tables for Figures 4(b-d) are Tables S13-S15 in Supplementary Additional file. For Dataset3, Tables S16-18 correspond to Figures 6(b-d) in Supplementary Additional file.

It is worth noting that Tables 2, 4, and 6 were populated based on results from an enrichment analysis tool, Enrichr. For a set of terms in a category, testing the null hypothesis can tell if a user’s gene list enriched in given term (i.e., overlap column) is more than a random chance (or not) compared to that term’s gene list in the human genome background [18]. The terms in a category are ranked by p-values, which were derived using Fisher’s exact test and the adjusted p-values were corrected based on Benjamini–Hochberg procedure [15]. In our study, we had two categories in Enrichr: ARCHS4 Tissues and NCI-60 Cancer Cell Lines, reporting one term (BREAST (BULK TISSUE)) in the former and three terms (MD-MB231, MCF7, HS578T) in the latter. These breast cancer cell lines are well-established in biology and medicine when analysing breast cancer as in [98–103].

## 5 Conclusion and Future Work

In this paper, we present a computational framework based on machine learning and enrichment analysis, unveiling critical genes, drugs, drug targets and other biological knowledge underlying breast cancer drug response mechanism. Our framework receives as an input a gene expression dataset pertaining to breast cancer patients responding and not responding to a treatment; we downloaded three different gene expression datasets from the GEO database according to the following accession numbers: GSE130787, GSE140494, and GSE196093. As 𝑛 ≫ 𝑚 arises challenge in computational genomics in which *n* and *m* are the number of genes and samples, respectively, we formulate our method according to the dual form in which we solve the optimization problem as a function of λ associated with *m* (instead of the formulation as a function of w associated with *n*). Then, we perform mathematical calculations to efficiently recover w as a function of three inputs (λ , y, x) and then identifying *p* important genes out of *n*, provided to enrichment analysis tools Enrichr and Metascape. In addition to (1) significantly achieving the highest performance results in terms of the area under curve; and (2) being more computationally efficient than the baseline SVM, results demonstrate that our method esvm outperformed existing baseline methods including deep learning in (1) breast cancer cell line identification, showing more expressed genes; and (2) achieving the highest performance results from a classification perspective when coupled with SVM. Moreover, we reported several drugs (including tamoxifen, cisplatin, and erlotinib), 36 unique TFs (e.g., SP1, NFKB1, RELA), and 74 unique genes (including tumor suppression genes such as TP53, BRCA1, and RB1) that have been reported to be connected to drug response and resistance mechanism, progression, and metastasis of breast cancer. We made our computational method available publicly on the maGENEgerZ web server at https://aibio.shinyapps.io/maGENEgerZ/.

Future work includes (1) utilizing our framework to unveil various biological knowledge behind drug response mechanisms, related to different cancer types such as pancreatic cancer, liver cancer, and multiple myeloma; (2) collaborating with clinical research physicians to apply our tool to analyze drug response mechanism in the neoadjuvant setting; and (3) integrating different profiling data related to cancer drug response and performing an assessment from biological perspective.

## CRediT authorship contribution statement

**Turki Turki:** Conceptualization, Formal analysis, Methodology, Data curation, Software, Supervision, Writing - original draft, Visualization, Investigation. **Y-h. Taguchi:** Conceptualization, Formal analysis, Methodology, Data curation, Validation, Writing - original draft.

**Turki Turki** received a B.S. in computer science from King Abdulaziz University, an M.S. in computer science from NYU.POLY, and a Ph.D. in computer science from the New Jersey Institute of Technology. He is currently an associate professor with the Department of Computer Science, King Abdulaziz University, Saudi Arabia. His research interests include machine learning, data science and bioinformatics. His research has been published in journals such as *Scientific Reports, IEEE Journal on Selected Topics in Signal Processing*, *Frontiers in Genetics and Computers in Biology and Medicine*. He is an editorial board member of *Sustainable Computing: Informatics and Systems, BMC Medical Genomics, PLOS ONE*, and *Computers in Biology and Medicine*.

**Y-h. Taguchi** received a B.S. degree in physics from the Tokyo Institute of Technology and a Ph.D. degree in physics from the Tokyo Institute of Technology. He is currently a full professor with the Department of Physics, Chuo University, Japan. His works have been published in leading journals such as *Physical Review Letters*, *Bioinformatics*, and *Scientific Reports*. His research interests include bioinformatics, machine learning, and nonlinear physics. He is also an editorial board member of *PloS ONE*, *BMC Medical Genomics*, *Medicine* (Lippincott Williams & Wilkins journal), *BMC Research Notes*, and *IPSJ Transaction on Bioinformatics*.

