## Supplementary material for "maGENEgerZ: An Efficient AI-Based Framework Can Extract More Expressed Genes and Biological Insights Underlying Breast Cancer Drug Response Mechanism": Additional File.docx

| Linear: | 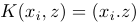 | (Eq. S1) |
| --- | --- | --- |
| Polynomial: | 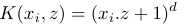 | (Eq. S2) |
| Gaussian: | 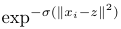 | (Eq. S3) |
| Sigmoid: | 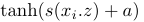 | (Eq. S4) |
| Laplacian: | 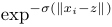 | (Eq. S5) |

### **Equations**

### **Tables**

**Table S1**: Enriched terms obtained from a) ARCHS4 Tissues and b) NCI-60 Cancer Cell Lines via Enrichr according to produced genes by each deep learning method when using Dataset1. The best result is shown in bold.

| **a) ARCHS4 Tissues** | | | | | |
| --- | --- | --- | --- | --- | --- |
| **Method** | **Rank** | **Term** | **Overlap** | ***P*-value** | **Adjusted *P*-value** |
| DeepLIFT | 2 | BREAST (BULK TISSUE) | 40/2316 | 3.06×10^-13^ | 1.65×10^-11^ |
| DeepSHAP | 6 | BREAST (BULK TISSUE) | 29/2316 | 1.87×10^-6^ | 3.36×10^-5^ |
| LRP | 3 | BREAST (BULK TISSUE) | 35/2316 | 7.13×10^-10^ | 2.56×10^-8^ |
| **b) NCI-60 Cancer Cell Lines** | | | | | |
| **Method** | **Rank** | **Term** | **Overlap** | ***P*-value** | **Adjusted *P*-value** |
| DeepLIFT | 42 | MD-MB231 | 1/150 | 5.29×10^-1^ | 9.28×10^-1^ |
|  | 27 | MCF7 | 3/397 | 3.19×10^-1^ | 8.01×10^-1^ |
|  | 18 | HS578T | 2/176 | 2.19×10^-1^ | 8.01×10^-1^ |
| DeepSHAP | - | MD-MB231 | - | - | - |
|  | 52 | MCF7 | 2/397 | 5.93×10^-1^ | 7.89×10^-1^ |
|  | - | HS578T | - | - | - |
| LRP | 44 | MD-MB231 | 1/150 | 5.29×10^-1^ | 8.44×10^-1^ |
|  | 69 | MCF7 | 1/397 | 8.66×10^-1^ | 9.38×10^-1^ |
|  | 17 | HS578T | 2/176 | 2.19×10^-1^ | 7.95×10^-1^ |

**Table S2**: Enriched terms obtained from a) ARCHS4 Tissues and b) NCI-60 Cancer Cell Lines via Enrichr according to produced genes by each deep learning method when using Dataset2. The best result is shown in bold.

| **a) ARCHS4 Tissues** | | | | | |
| --- | --- | --- | --- | --- | --- |
| **Method** | **Rank** | **Term** | **Overlap** | ***P*-value** | **Adjusted *P*-value** |
| DeepLIFT | 2 | BREAST (BULK TISSUE) | 21/2316 | 4.75×10^-3^ | 1.69×10^-1^ |
| DeepSHAP | 22 | BREAST (BULK TISSUE) | 17/2316 | 6.70×10^-2^ | 2.68×10^-1^ |
| LRP | 34 | BREAST (BULK TISSUE) | 7/2316 | 9.52×10^-1^ | 9.99×10^-1^ |
| **b) NCI-60 Cancer Cell Lines** | | | | | |
| **Method** | **Rank** | **Term** | **Overlap** | ***P*-value** | **Adjusted *P*-value** |
| DeepLIFT | 57 | MD-MB231 | 1/150 | 5.29×10^-1^ | 7.59×10^-1^ |
|  | 63 | MCF7 | 2/397 | 5.93×10^-1^ | 7.81×10^-1^ |
|  | 30 | HS578T | 2/176 | 2.19×10^-1^ | 5.74×10^-1^ |
| DeepSHAP | - | MD-MB231 | - | - | - |
|  | 16 | MCF7 | 4/397 | 1.37×10^-1^ | 6.98×10^-1^ |
|  | 47 | HS578T | 1/176 | 5.87×10^-1^ | 9.07×10^-1^ |
| LRP | - | MD-MB231 | - | - | - |
|  | 39 | MCF7 | 3/397 | 3.19×10^-1^ | 6.35×10^-1^ |
|  | 10 | HS578T | 3/176 | 5.83×10^-2^ | 4.47×10^-1^ |

**Table S3**: Enriched terms obtained from a) ARCHS4 Tissues and b) NCI-60 Cancer Cell Lines via Enrichr according to produced genes by each deep learning method when using Dataset3. The best result is shown in bold.

| **a) ARCHS4 Tissues** | | | | | |
| --- | --- | --- | --- | --- | --- |
| **Method** | **Rank** | **Term** | **Overlap** | ***P*-value** | **Adjusted *P*-value** |
| DeepLIFT | 9 | BREAST (BULK TISSUE) | 7/2316 | 3.57×10^-1^ | 9.97×10^-1^ |
| DeepSHAP | 2 | BREAST (BULK TISSUE) | 7/2316 | 3.57×10^-1^ | 9.97×10^-1^ |
| LRP | 3 | BREAST (BULK TISSUE) | 6/2316 | 5.28×10^-1^ | 9.97×10^-1^ |
| **b) NCI-60 Cancer Cell Lines** | | | | | |
| **Method** | **Rank** | **Term** | **Overlap** | ***P*-value** | **Adjusted *P*-value** |
| DeepLIFT | - | MD-MB231 | **-** | - | - |
|  | 23 | MCF7 | 2/397 | 2.61×10^-1^ | 6.10×10^-1^ |
|  | - | HS578T | **-** | - | - |
| DeepSHAP | - | MD-MB231 | - | - | - |
|  | 18 | MCF7 | 2/397 | 2.61×10^-1^ | 6.12×10^-1^ |
|  | - | HS578T | - | - | - |
| LRP | - | MD-MB231 | - | - | - |
|  | 20 | MCF7 | 2/397 | 2.61×10^-1^ | 6.42×10^-1^ |
|  | - | HS578T | **-** | - | - |

| **Model** | **ACC** | **BAC** | **F1** | **MCC** |
| --- | --- | --- | --- | --- |
| esvm | **0.921**(0.076) | **0.935**(0.059) | **0.926**(0.067) | **0.849(0.147)** |
| limma | 0.529(0.071) | 0.547(0.076) | 0.583(0.092) | 0.080(0.136) |
| sam | 0.521(0.126) | 0.544(0.118) | 0.581(0.128) | 0.087(0.247) |
| *t*-test | 0.541(0.080) | 0.558(0.091) | 0.604(0.112) | 0.110(0.169) |
| lasso | 0.495(0.090) | 0.505(0.109) | 0.557(0.099) | 0.001(0.196) |
| DeepLIFT | 0.526(0.124) | 0.482(0.161) | 0.582(0.091) | -0.023(0.307) |
| DeepSHAP | 0.596(0.090) | 0.600(0.088) | 0.643(0.114) | 0.188(0.155) |
| LRP | 0.482(0.052) | 0.455(0.060) | 0.550(0.117) | -0.084(0.121) |
| None | 0.642(0.124) | 0.611(0.121) | 0.709(0.115) | 0.243(0.266) |

**Table S4***:* Generalization (testing) performance results (and standard deviation) on Dataset1 when utilizing five-fold cross-validation according to SVM coupled with obtained genes via each method. ACC is accuracy. BAC is balanced accuracy. MCC is Matthews correlation coefficient. None refers to the complete set of genes in the original dataset.

**Table S5***:* Generalization (testing) performance results (and standard deviation) on Dataset2 when utilizing five-fold-cross-validation according to SVM coupled with obtained genes via each method. ACC is accuracy. BAC is balanced accuracy. MCC is Matthews correlation coefficient. None refers to the complete set of genes in the original dataset.

| **Model** | **ACC** | **BAC** | **F1** | **MCC** |
| --- | --- | --- | --- | --- |
| esvm | **0.978**(0.047) | **0.960**(0.089) | **0.950**(0.111) | **0.944**(0.123) |
| limma | 0.691(0.032) | 0.522(0.087) | 0.246(0.159) | 0.069(0.165) |
| sam | 0.734(0.075) | 0.579(0.055) | 0.303(0.063) | 0.168(0.115) |
| *t*-test | 0.656(0.153) | 0.549(0.104) | 0.266(0.179) | NA |
| lasso | 0.635(0.133) | 0.481(0.087) | 0.210(0.148) | -0.033(0.162) |
| DeepLIFT | 0.723(0.064) | 0.565(0.111) | 0.290(0.181) | 0.180(0.218) |
| DeepSHAP | 0.666(0.093) | 0.511(0.097) | 0.166(0.155) | -0.015(0.206) |
| LRP | 0.644(0.098) | 0.542(0.104) | 0.243(0.171) | 0.046(0.191) |
| None | 0.787(0.113) | 0.541(0.117) | 0.133(0.298) | NA |

**Table S6***:* Generalization (testing) performance results (and standard deviation) on Dataset3 when utilizing five-fold cross-validation according to SVM coupled with obtained genes via each method. ACC is accuracy. BAC is balanced accuracy. MCC is Matthews correlation coefficient. None refers to the complete set of genes in the original dataset.

| **Model** | **ACC** | **BAC** | **F1** | **MCC** |
| --- | --- | --- | --- | --- |
| esvm | **0.726**(0.028) | **0.676**(0.046) | **0.801**(0.019) | **0.377**(0.083) |
| limma | 0.660(0.006) | 0.528(0.016) | 0.787(0.002) | 0.117(0.055) |
| sam | 0.692(0.022) | 0.605(0.028) | 0.791(0.016) | 0.266(0.065) |
| *t*-test | 0.692(0.022) | 0.605(0.028) | 0.791(0.016) | 0.266(0.065) |
| lasso | 0.676(0.011) | 0.555(0.015) | 0.793(0.008) | 0.197(0.064) |
| DeepLIFT | 0.680(0.038) | 0.615(0.037) | 0.771(0.032) | 0.258(0.086) |
| DeepSHAP | 0.668(0.015) | 0.604(0.017) | 0.761(0.025) | 0.231(0.024) |
| LRP | 0.663(0.024) | 0.606(0.029) | 0.753(0.025) | 0.227(0.060) |
| None | 0.642(0.046) | 0.604(0.054) | 0.725(0.047) | 0.213(0.109) |

**Table S7***:* Generalization (testing) performance results (and standard deviation) on Dataset1 when utilizing five-fold cross-validation according to lasso coupled with obtained genes via each method. ACC is accuracy. BAC is balanced accuracy. MCC is Matthews correlation coefficient. None refers to the complete set of genes in the original dataset.

| **Model** | **ACC** | **BAC** | **F1** | **MCC** |
| --- | --- | --- | --- | --- |
| esvm | **0.662**(0.104) | **0.660**(0.101) | **0.738**(0.080) | **NA** |
| limma | 0.607(0.082) | 0.593(0.060) | 0.688(0.096) | NA |
| sam | 0.543(0.136) | 0.482(0.025) | 0.677(0.126) | NA |
| *t*-test | 0.554(0.139) | 0.514(0.040) | 0.699(0.106) | NA |
| lasso | 0.564(0.178) | 0.484(0.064) | 0.702(0.153) | NA |
| DeepLift | 0.520(0.142) | 0.456(0.067) | 0.675(0.119) | NA |
| DeepSHAP | 0.576(0.158) | 0.500(0.000) | 0.721(0.126) | NA |
| LRP | 0.509(0.151) | 0.453(0.103) | 0.665(0.122) | NA |
| None | 0.589(0.188) | 0.544(0.111) | 0.686(0.168) | NA |

**Table S8***:* Generalization (testing) performance results (and standard deviation) on Dataset2 when utilizing five-fold cross-validation according to lasso coupled with obtained genes via each method. ACC is accuracy. BAC is balanced accuracy. MCC is Matthews correlation coefficient. None refers to the complete set of genes in the original dataset.

| **Model** | **ACC** | **BAC** | **F1** | **MCC** |
| --- | --- | --- | --- | --- |
| esvm | 0.787(0.098) | **0.537**(0.080) | 0.157(0.228) | NA |
| limma | **0.788**(0.077) | 0.500(0.000) | NA | NA |
| sam | **0.788**(0.077) | 0.500(0.000) | NA | NA |
| *t*-test | **0.788**(0.077) | 0.500(0.000) | NA | NA |
| lasso | **0.788**(0.077) | 0.500(0.000) | NA | NA |
| DeepLIFT | **0.788**(0.077) | 0.500(0.000) | NA | NA |
| DeepSHAP | **0.788**(0.077) | 0.500(0.000) | NA | NA |
| LRP | **0.788**(0.077) | 0.500(0.000) | NA | NA |
| None | **0.788**(0.077) | 0.500(0.000) | NA | NA |

**Table S9***:* Generalization (testing) performance results (and standard deviation) on Dataset3 when utilizing five-fold cross-validation according to lasso coupled with obtained genes via each method. ACC is accuracy. BAC is balanced accuracy. MCC is Matthews correlation coefficient. None refers to the complete set of genes in the original dataset.

| **Model** | **ACC** | **BAC** | **F1** | **MCC** |
| --- | --- | --- | --- | --- |
| esvm | 0.690(0.020) | **0.627**(0.035) | 0.778(0.015) | **0.280**(0.067) |
| limma | 0.649(0.004) | 0.513(0.011) | 0.781(0.007) | NA |
| sam | **0.697**(0.023) | 0.613(0.035) | 0.792(0.015) | 0.277(0.079) |
| *t*-test | 0.694(0.023) | 0.610(0.035) | 0.791(0.014) | 0.269(0.075) |
| lasso | 0.676(0.018) | 0.563(0.013) | 0.790(0.014) | 0.199(0.047) |
| DeepLIFT | **0.697**(0.037) | 0.605(0.047) | **0.796**(0.024) | 0.272(0.115) |
| DeepSHAP | 0.694(0.026) | 0.607(0.033) | 0.792(0.018) | 0.269(0.079) |
| LRP | 0.695(0.020) | 0.615(0.035) | 0.790(0.013) | 0.274(0.069) |
| None | 0.687(0.024) | 0.588(0.028) | 0.792(0.017) | 0.239(0.081) |

**Table S10:** Forty-four genes from PPI based on genes of esvm in Dataset1 coupled with Metascape.

**GRP TRH LTF TTR APOD CSSC4 CAPN9 CAPN13 S100A6 S100A2 CARTPT DUSP13B FKBP10 GGH ERP27 POF1B SOSTDC1 CEACAM5 BMP6 CHRDL2 GSTA1 QDPR NDRG1 NR4A1 SOCs4 PPBP CCL21 APLN TRIM63 ADCY5 CALML4 MYO5C RAB38 AQP3 TUBA3C PTP4A3 FOS MS4A2 MMP1 MMP3 MMP12 FGFR2 IGF1 PGR**

**Table S11:** Transcription factors (TFs) according to Metascape when coupled with genes from esvm in Dataset1.

| **TF** | -**log10(P)** |
| --- | --- |
| NFKBIA | 4.4 |
| FOS | 4.2 |
| RELA | 3.9 |
| STAT3 | 3.7 |
| JUN | 3.6 |
| BRCA1 | 3.0 |
| SP1 | 2.7 |
| USF1 | 2.7 |
| ETS1 | 2.6 |
| CREB1 | 2.3 |
| NFKB1 | 2.3 |
| MYC | 2.2 |

| **Term** | -**log10(P)** |
| --- | --- |
| WP2877: Vitamin D receptor pathway | 6.683087629 |
| M5885: NABA MATRISOME ASSOCIATED | 5.929437687 |
| GO:0030879: mammary gland development | 5.461914176 |
| R-HSA-1474244: Extracellular matrix organization | 5.148258272 |
| GO:1903530: regulation of secretion by cell | 4.730109472 |
| P06664: Gonadotropin-releasing hormone receptor pathway | 4.722968047 |
| GO:0060349: bone morphogenesis | 4.682426759 |
| GO:0002067: glandular epithelial cell differentiation | 4.188738536 |
| GO:0071396: cellular response to lipid | 4.171174811 |
| WP3942: PPAR signaling pathway | 4.111813107 |
| GO:0071456: cellular response to hypoxia | 4.089391086 |
| GO:0035239: tube morphogenesis | 4.073369608 |
| hsa04024: cAMP signaling pathway | 3.981041666 |
| R-HSA-287179: FCERI mediated MAPK activation | 3.792331362 |
| GO:0051051: negative regulation of transport | 3.778828799 |
| GO:0045576: mast cell activation | 3.751978524 |
| WP2840: Hair follicle development: cytodifferentiation - part 3 of 3 | 3.695955709 |
| GO:0006576: biogenic amine metabolic process | 3.695955709 |
| GO:0007631: feeding behavior | 3.657970614 |
| GO:0031349: positive regulation of defense response | 3.330329045 |

**Table S12:** Process and pathway enrichment analysis provided by Metascape according to genes from esvm using Dataset1.

**Table S13:** Twelve genes from PPI based on genes of esvm in Dataset2 coupled with Metascape.

**ACTG2 PRKAR2B ALDH2 PRKACB KRT19 KRT18 WASF3 DLG5 TRIM37 GSTM3 GSTA1 CYP2A6**

**Table S14:** Transcription factors (TFs) according to Metascape when coupled with genes from esvm in Dataset2.

| **TF** | -**log10(P)** |
| --- | --- |
| SP1 | 14 |
| RELA | 7.8 |
| NFKB1 | 7.8 |
| HIF1A | 6.0 |
| STAT3 | 4.8 |
| ZEB1 | 4.6 |
| FOXO3 | 4.4 |
| NFE2L2 | 4.4 |
| JUN | 3.6 |
| NR3C1 | 3.1 |
| SP3 | 3.1 |
| EP300 | 2.9 |
| E2F1 | 2.9 |
| CEBPB | 2.8 |
| USF1 | 2.7 |
| HDAC1 | 2.6 |
| ETS1 | 2.6 |
| STAT1 | 2.4 |

**Table S15:** Process and pathway enrichment analysis provided by Metascape according to genes from esvm using Dataset2.

| **Term** | -**log10(P)** |
| --- | --- |
| WP2882: Nuclear receptors meta-pathway | 7.027044378 |
| GO:0040008: regulation of growth | 6.900041718 |
| GO:0032119: sequestering of zinc ion | 6.856579571 |
| hsa05200: Pathways in cancer | 5.824415790 |
| GO:0010038: response to metal ion | 5.513319596 |
| R-HSA-1280218: Adaptive Immune System | 5.049127292 |
| R-HSA-211859: Biological oxidations | 5.033611143 |
| R-HSA-6785807: Interleukin-4 and Interleukin-13 signaling | 4.521040879 |
| hsa05202: Transcriptional misregulation in cancer | 4.348900537 |
| GO:0071900: regulation of protein serine/threonine kinase activity | 4.338211076 |
| GO:0097006: regulation of plasma lipoprotein particle levels | 4.325772820 |
| R-HSA-2022090: Assembly of collagen fibrils and other multimeric structures | 4.297415057 |
| GO:0009725: response to hormone | 4.296191723 |
| WP5094: Orexin receptor pathway | 4.250897679 |
| GO:0042445: hormone metabolic process | 4.191817439 |
| hsa04927: Cortisol synthesis and secretion | 4.188738536 |
| GO:0006656: phosphatidylcholine biosynthetic process | 4.118800000 |
| WP2880: Glucocorticoid receptor pathway | 4.062516151 |
| R-HSA-9759194: Nuclear events mediated by NFE2L2 | 3.857884409 |
| R-HSA-453279: Mitotic G1 phase and G1/S transition | 3.857191349 |

**Table S16:** Nineteen genes from PPI based on genes of esvm in Dataset3 coupled with Metascape.

**AR BIRC5 CCND1 RB1 STAT1 STAT3 ESR1 IRS1 PTEN ERBB2 ERBB3 ALK AKT1 MET JAK2 IGFIR EGFR TP53 MTOR**

**Table S17:** Transcription factors (TFs) according to Metascape when coupled with genes from esvm in Dataset3.

| **TF** | -**log10(P)** |
| --- | --- |
| TP53 | 25 |
| BRCA1 | 15 |
| HDAC1 | 14 |
| RELA | 14 |
| STAT3 | 13 |
| E2F1 | 12 |
| SP1 | 12 |
| NFKB1 | 10 |
| YBX1 | 9.5 |
| ESR1 | 9.5 |
| NKX3-1 | 9.2 |
| AR | 8.9 |
| VHL | 8.1 |
| PAX5 | 8.0 |
| CTNNB1 | 8.0 |
| PPARG | 7.8 |
| PGR | 7.7 |
| JUN | 7.7 |
| KDM4B | 7.5 |
| DNMT1 | 7.4 |

**Table S18:** Process and pathway enrichment analysis provided by Metascape according to genes from esvm using Dataset3.

| **Term** | -**log10(P)** |
| --- | --- |
| hsa05200: Pathways in cancer | 29.47897995 |
| WP4806: EGFR tyrosine kinase inhibitor resistance | 25.37029623 |
| WP2263: Androgen receptor network in prostate cancer | 23.74948849 |
| WP2034: Leptin signaling pathway | 23.47156603 |
| GO:0016310: phosphorylation | 19.38243190 |
| WP5087: Pleural mesothelioma | 17.49609721 |
| WP4172: PI3K-Akt signaling pathway | 17.23106359 |
| GO:0048732: gland development | 12.67768965 |
| WP3651: Pathways affected in adenoid cystic carcinoma | 12.42262917 |
| WP4685: Melanoma | 12.27973898 |
| WP3303: RAC1/PAK1/p38/MMP2 pathway | 12.27973898 |
| WP3850: Factors and pathways affecting insulin-like growth factor (IGF1)-Akt signaling | 11.88408602 |
| M91: PID TCPTP PATHWAY | 11.45566248 |
| WP138: Androgen receptor signaling pathway | 11.36503411 |
| GO:0030335: positive regulation of cell migration | 11.20280556 |
| GO:0040008: regulation of growth | 10.88191231 |
| WP4205: MET in type 1 papillary renal cell carcinoma | 10.52693300 |
| hsa04210: Apoptosis | 10.12258672 |
| hsa01524: Platinum drug resistance | 9.953967937 |
| R-HSA-9009391: Extra-nuclear estrogen signaling | 9.811315283 |

### **Figures**

| Dataset1  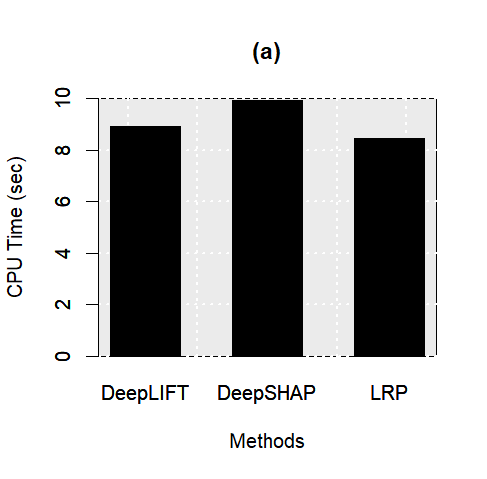 | Dataset2  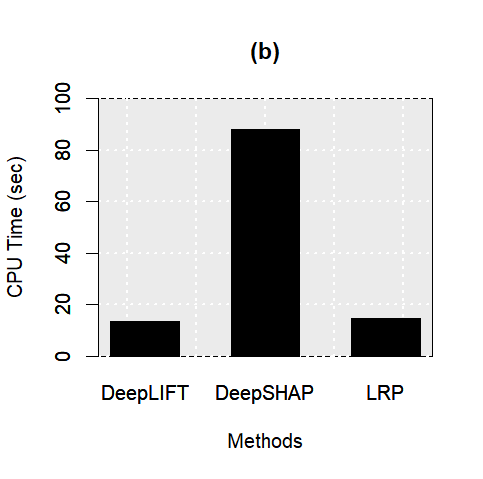 | Dataset3  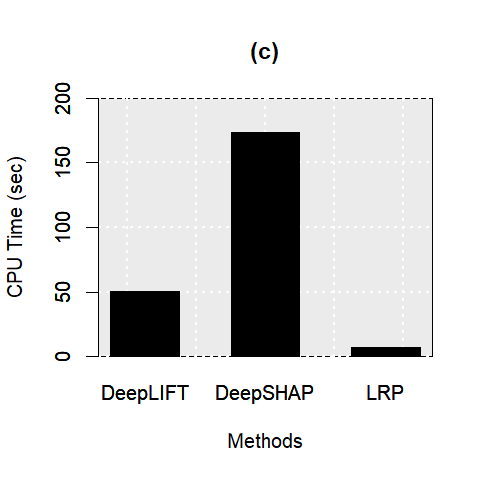 |
| --- | --- | --- |
| **Figure S1**: Computational running time for three deep learning methods based on three studied datasets. | | |

| **Dataset1** | DeepLIFT  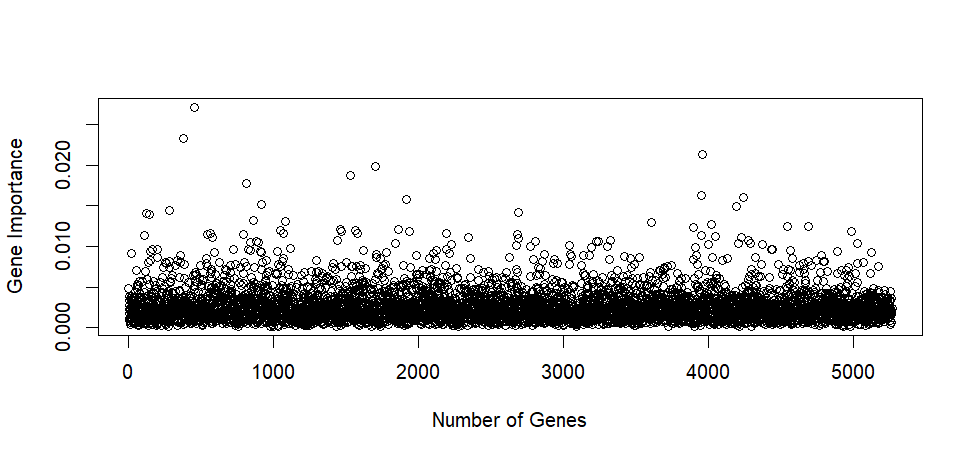 | DeepSHAP  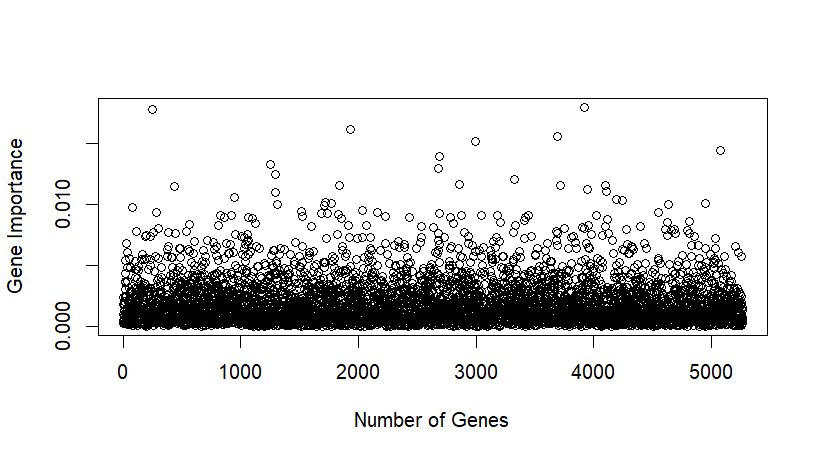 | LRP  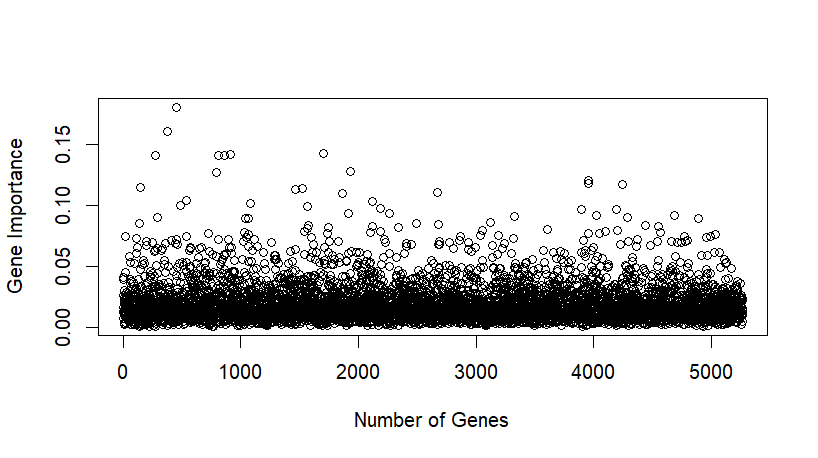 |
| --- | --- | --- | --- |
| **Dataset2** | DeepLIFT  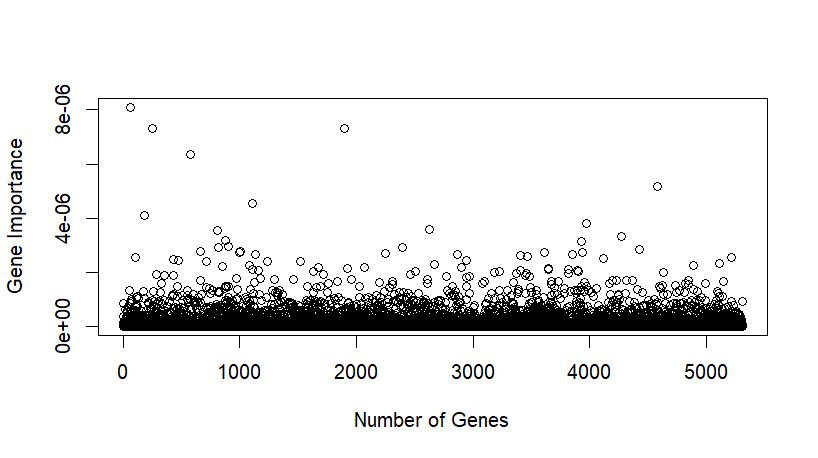 | DeepSHAP  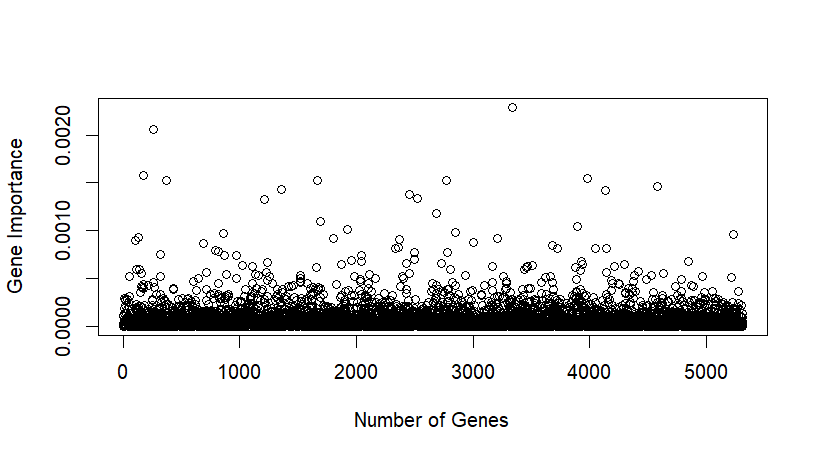 | LRP  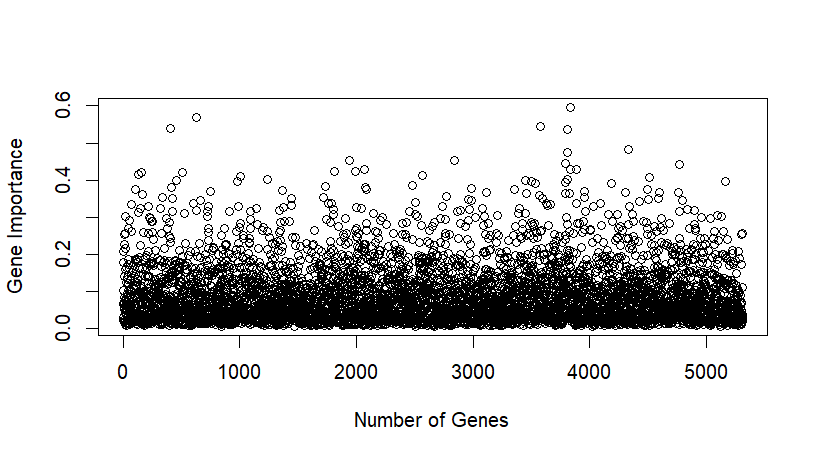 |
| **Dataset3** | DeepLIFT  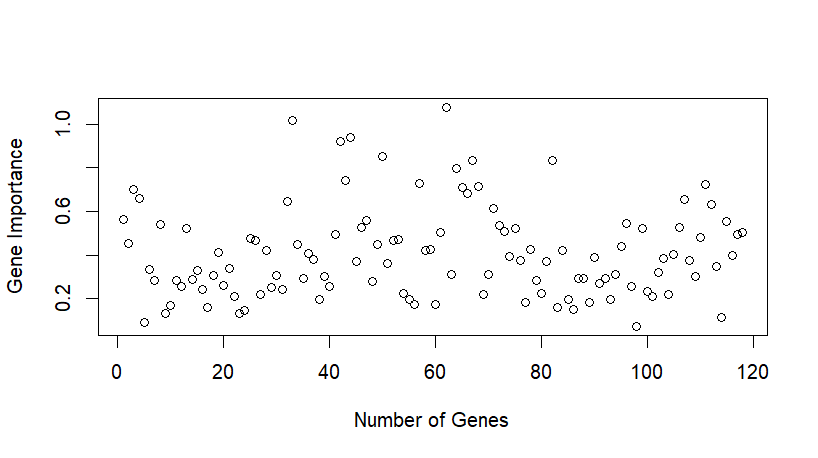 | DeepSHAP  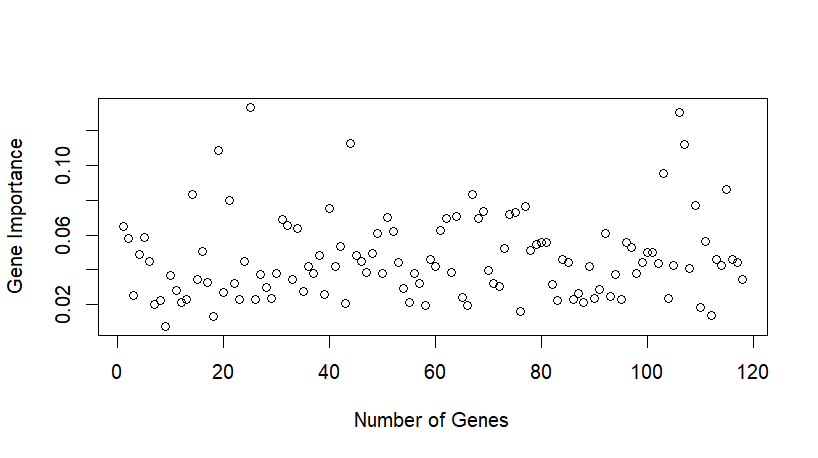 | LRP  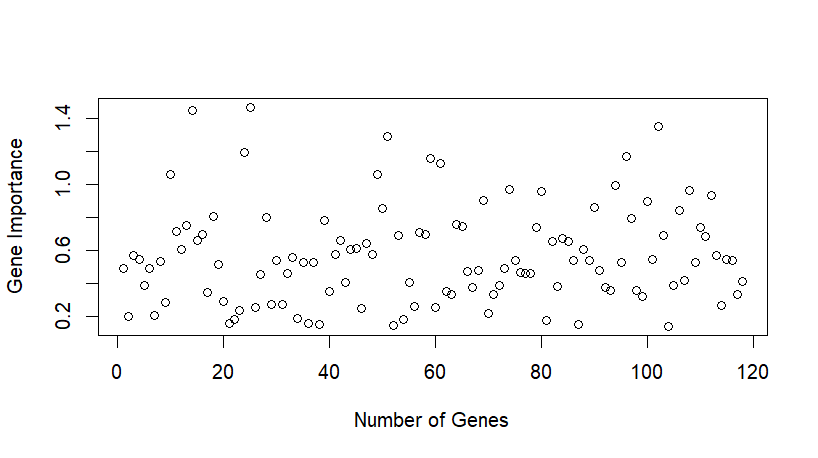 |
| **Figure S2**: Gene importance for three deep learning methods across the three studied datasets. | | | |

| 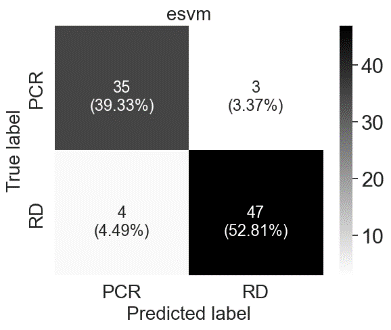 | 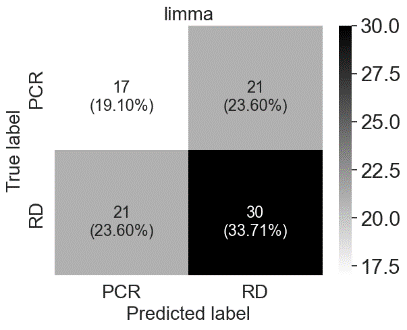 | 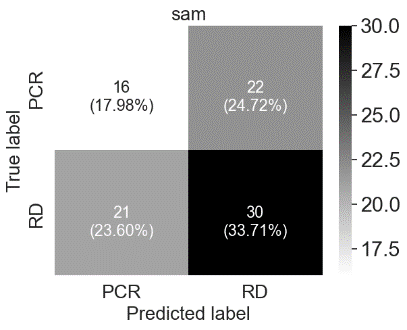 |
| --- | --- | --- |
| 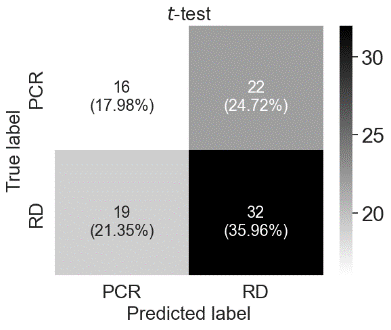 | 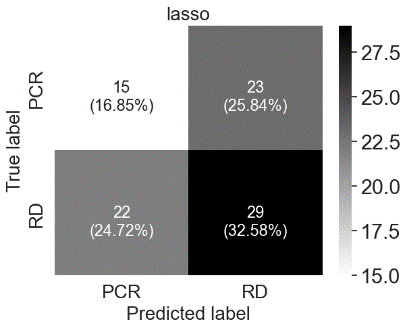 | 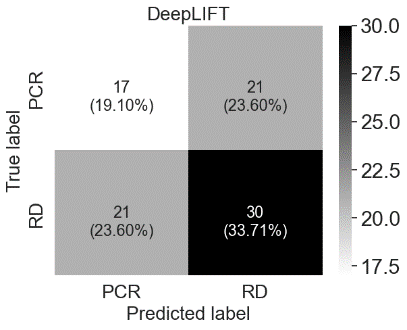 |
| 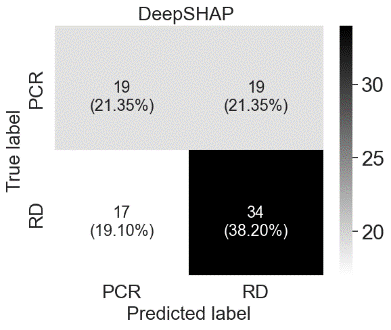 | 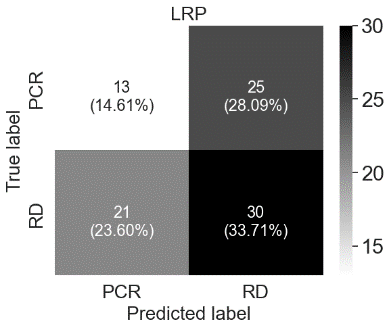 | 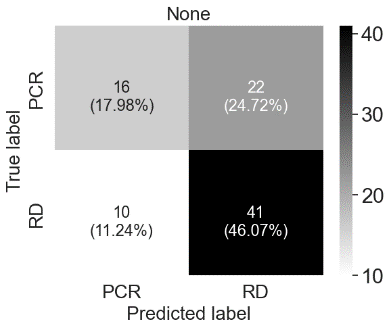 |
| **Figure S3:** Combined confusion matrices when running five-fold cross-validation on Dataset1 based on SVM coupled with genes provided via each method. None refers to the complete set of genes in the original dataset. | | |

| 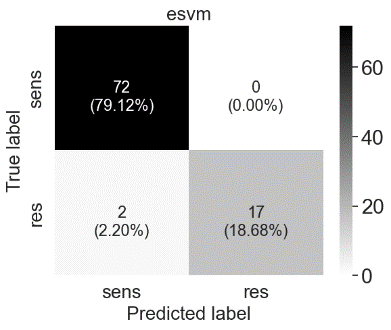 | 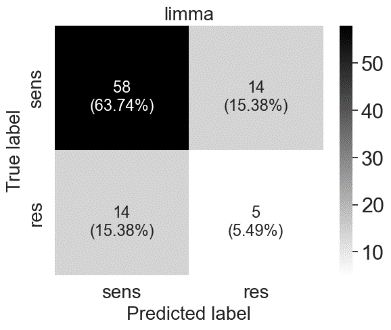 | 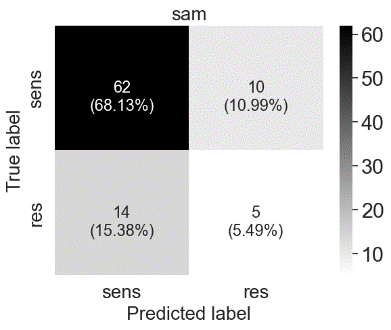 |
| --- | --- | --- |
| 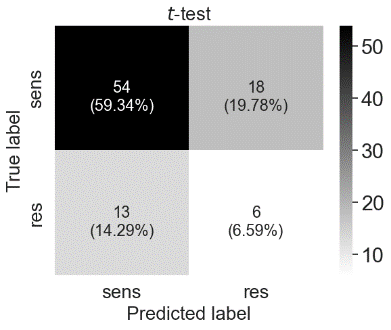 | 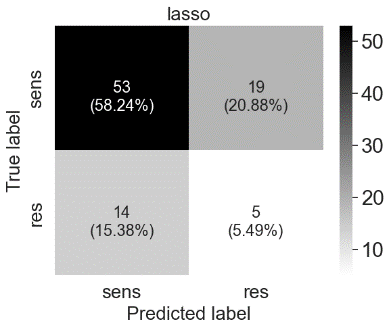 | 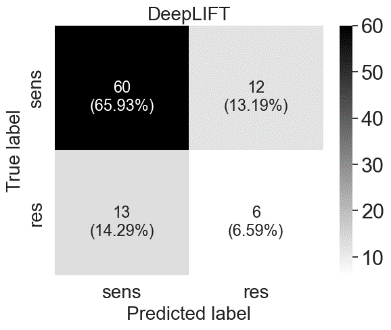 |
| 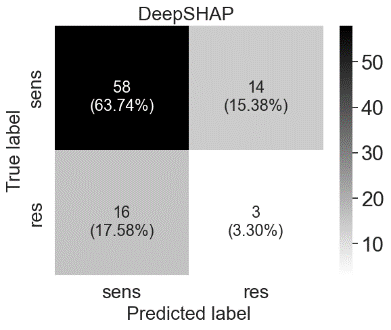 | 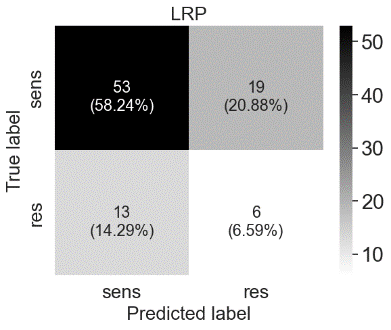 | 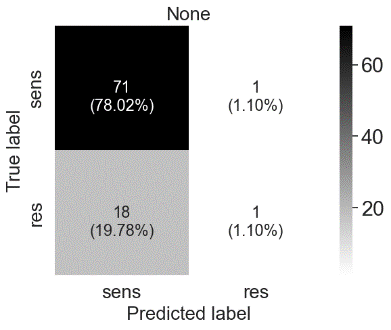 |
| **Figure S4:** Combined confusion matrices when running five-fold cross-validation on Dataset2 based on SVM coupled with genes provided via each method. None refers to the complete set of genes in the original dataset. | | |

| 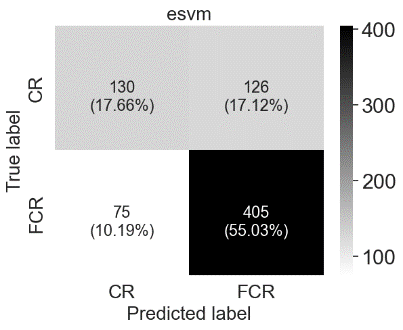 | 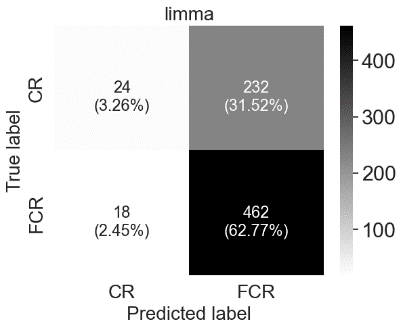 | 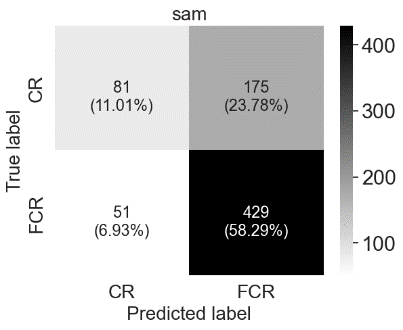 |
| --- | --- | --- |
| 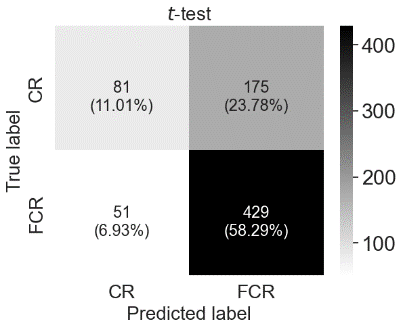 | 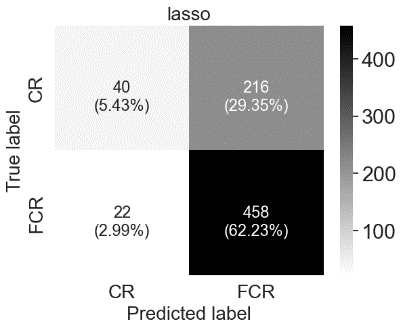 | 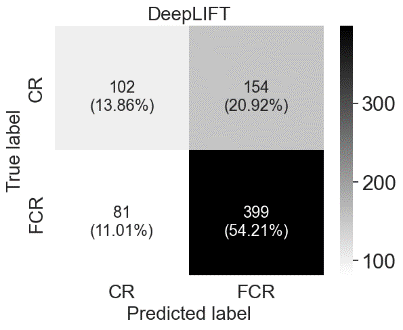 |
| 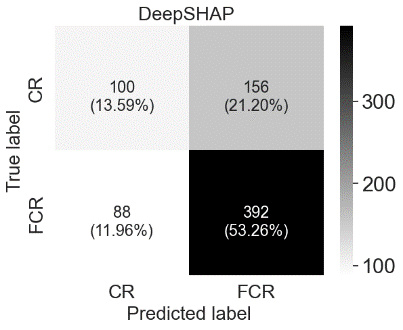 | 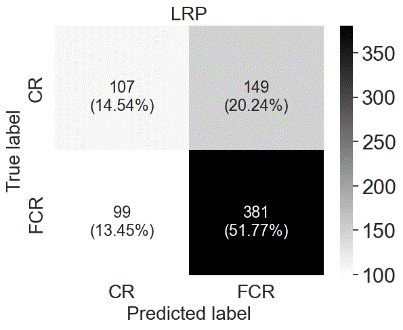 | 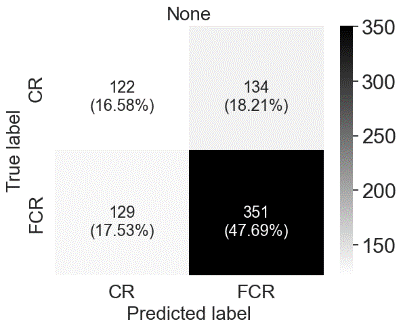 |
| **Figure S5:** Combined confusion matrices when running five-fold cross-validation on Dataset3 based on SVM coupled with genes provided via each method. None refers to the complete set of genes in the original dataset. | | |

| 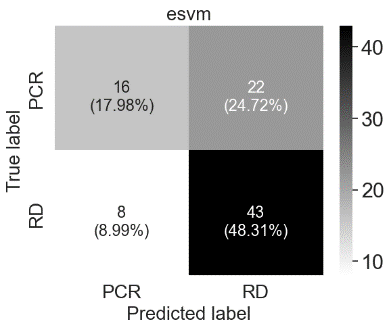 | 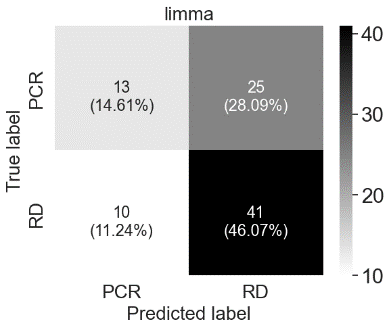 | 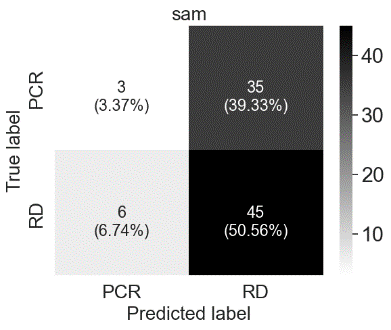 |
| --- | --- | --- |
| 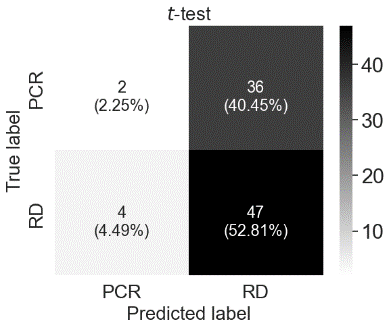 | 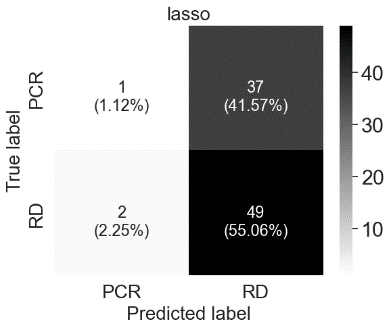 | 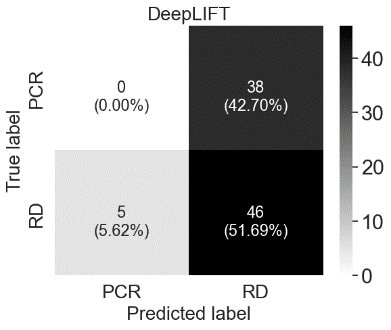 |
| 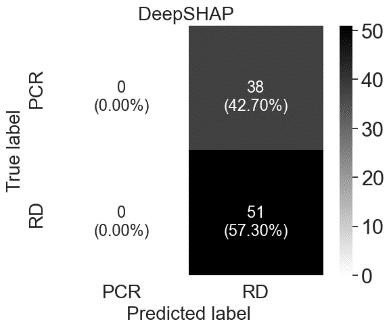 | 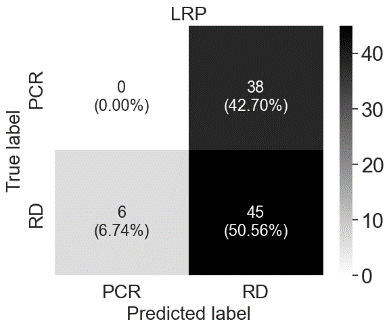 | 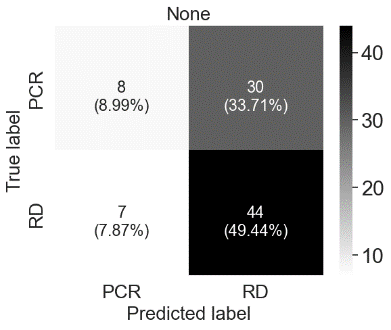 |
| **Figure S6:** Combined confusion matrices when running five-fold cross-validation on Dataset1 based on lasso coupled with genes provided via each method. None refers to the complete set of genes in the original dataset. | | |

| 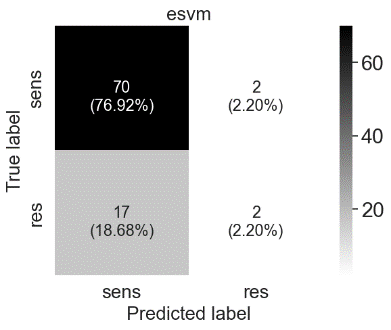 | 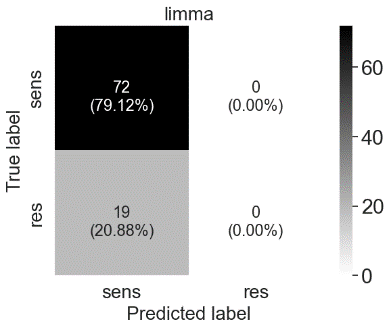 | 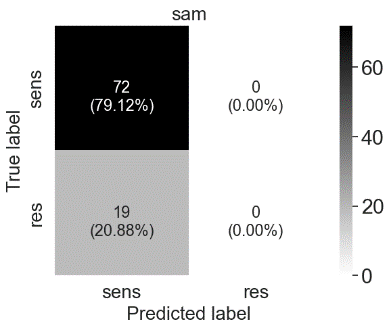 |
| --- | --- | --- |
| 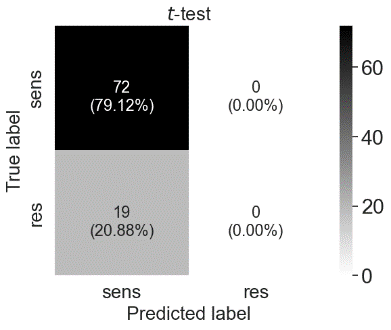 | 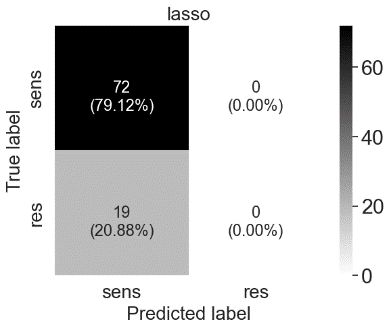 | 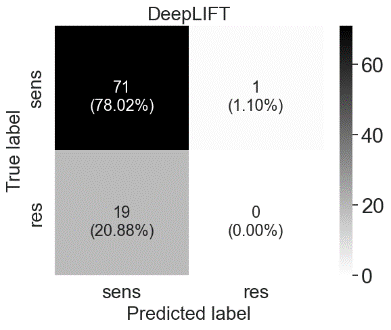 |
| 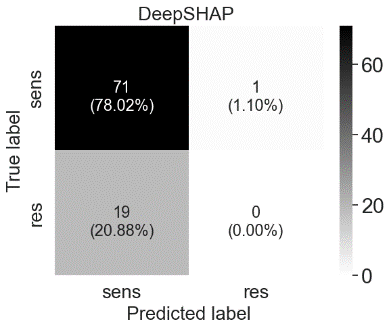 | 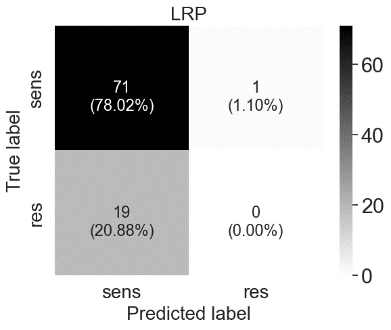 | 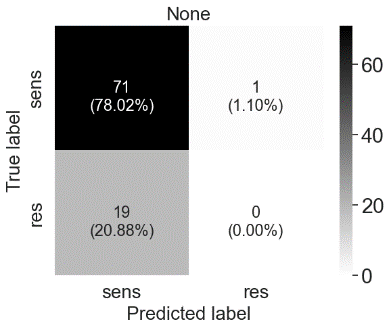 |
| **Figure S7:** Combined confusion matrices when running five-fold cross-validation on Dataset2 based on lasso coupled with genes provided via each method. None refers to the complete set of genes in the original dataset. | | |

| 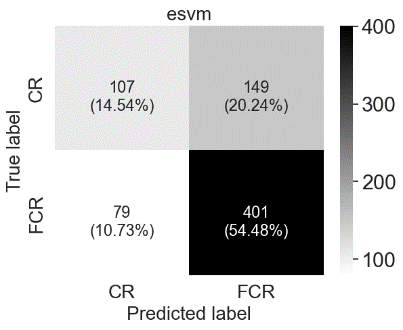 | 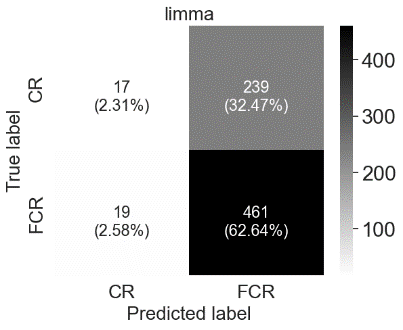 | 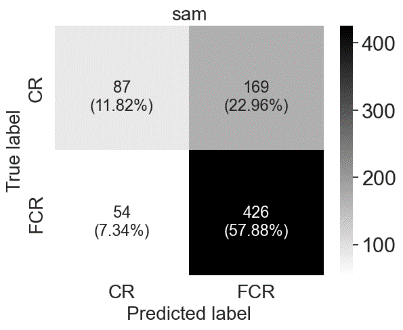 |
| --- | --- | --- |
| 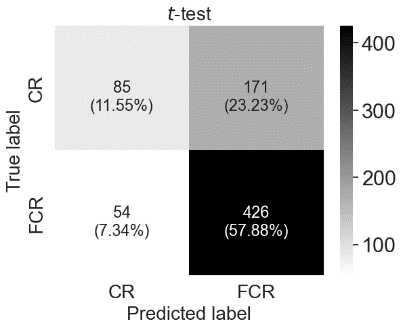 | 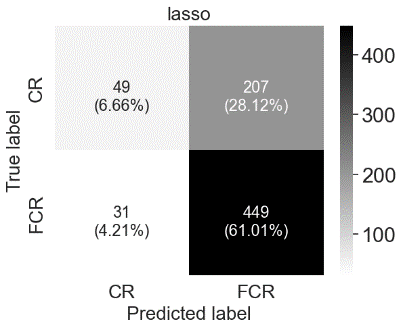 | 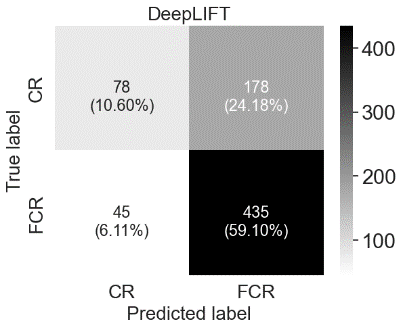 |
| 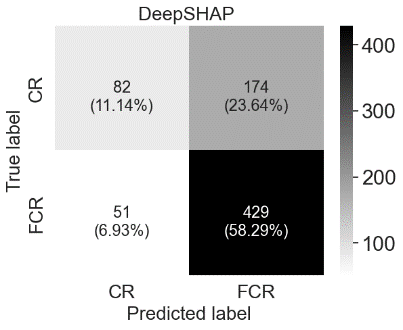 | 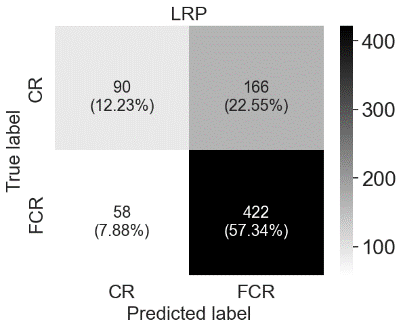 | 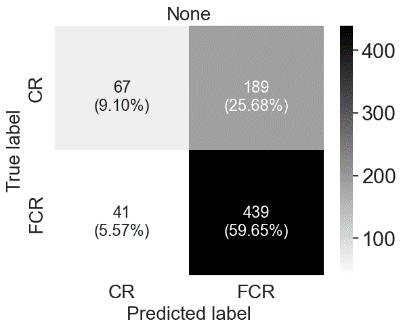 |
| **Figure S8:** Combined confusion matrices when running five-fold cross-validation on Dataset3 based on lasso coupled with genes provided via each method. None refers to the complete set of genes in the original dataset. | | |
