## Supplementary material for "maGENEgerZ: An Efficient AI-Based Framework Can Extract More Expressed Genes and Biological Insights Underlying Breast Cancer Drug Response Mechanism": maGENEgerZ_Screenshots.docx

- Go to <https://aibio.shinyapps.io/maGENEgerZ/>
- Upload the dataset from Datasets folder within Supplementary Materials. Suppose we upload Dataset1 named “GSE130787”. Then, we have the following.

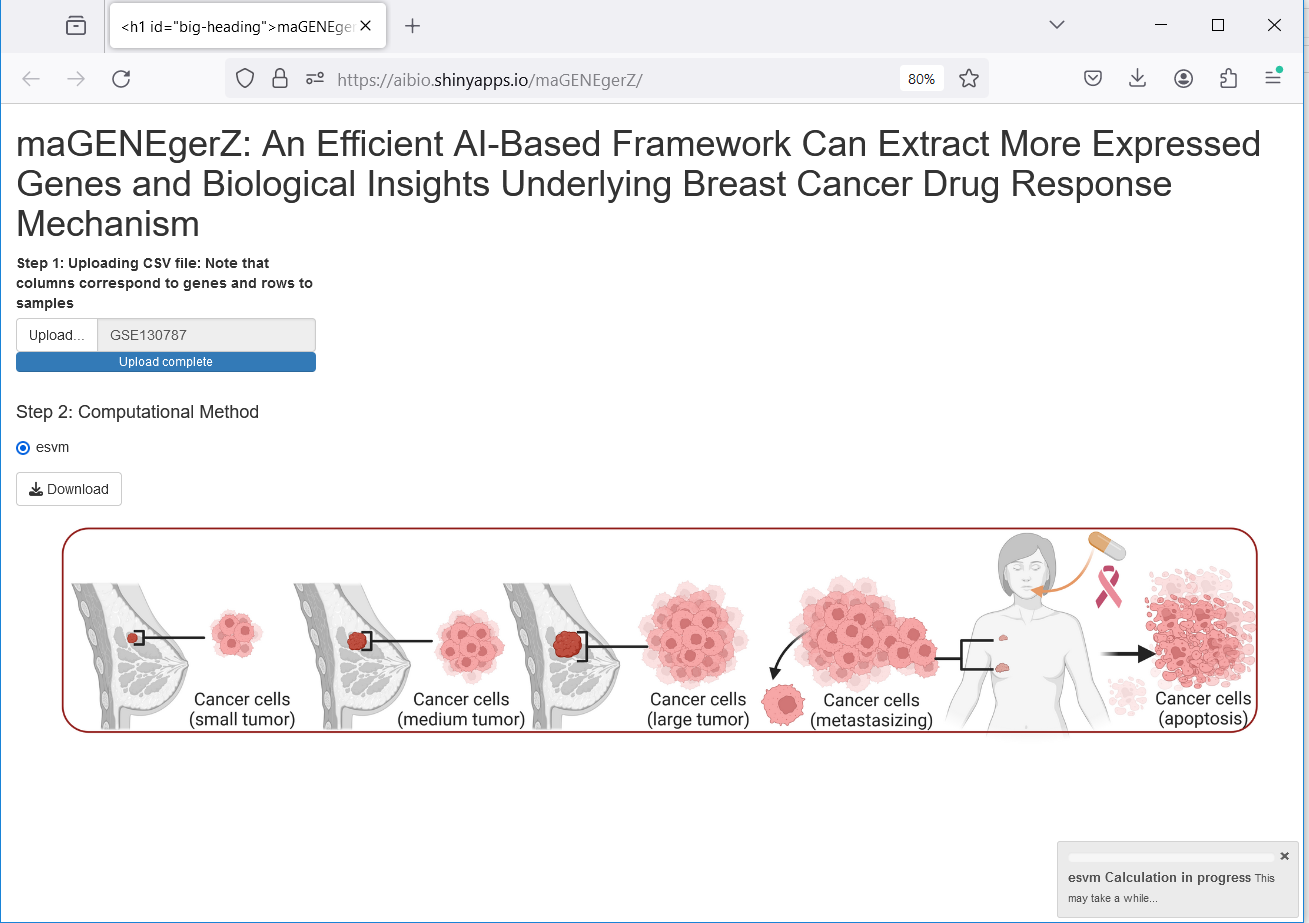

By default, esvm is selected and a progress bar showing the computation status.

After completion, we click on download button to download the results, shown as in the following screen.

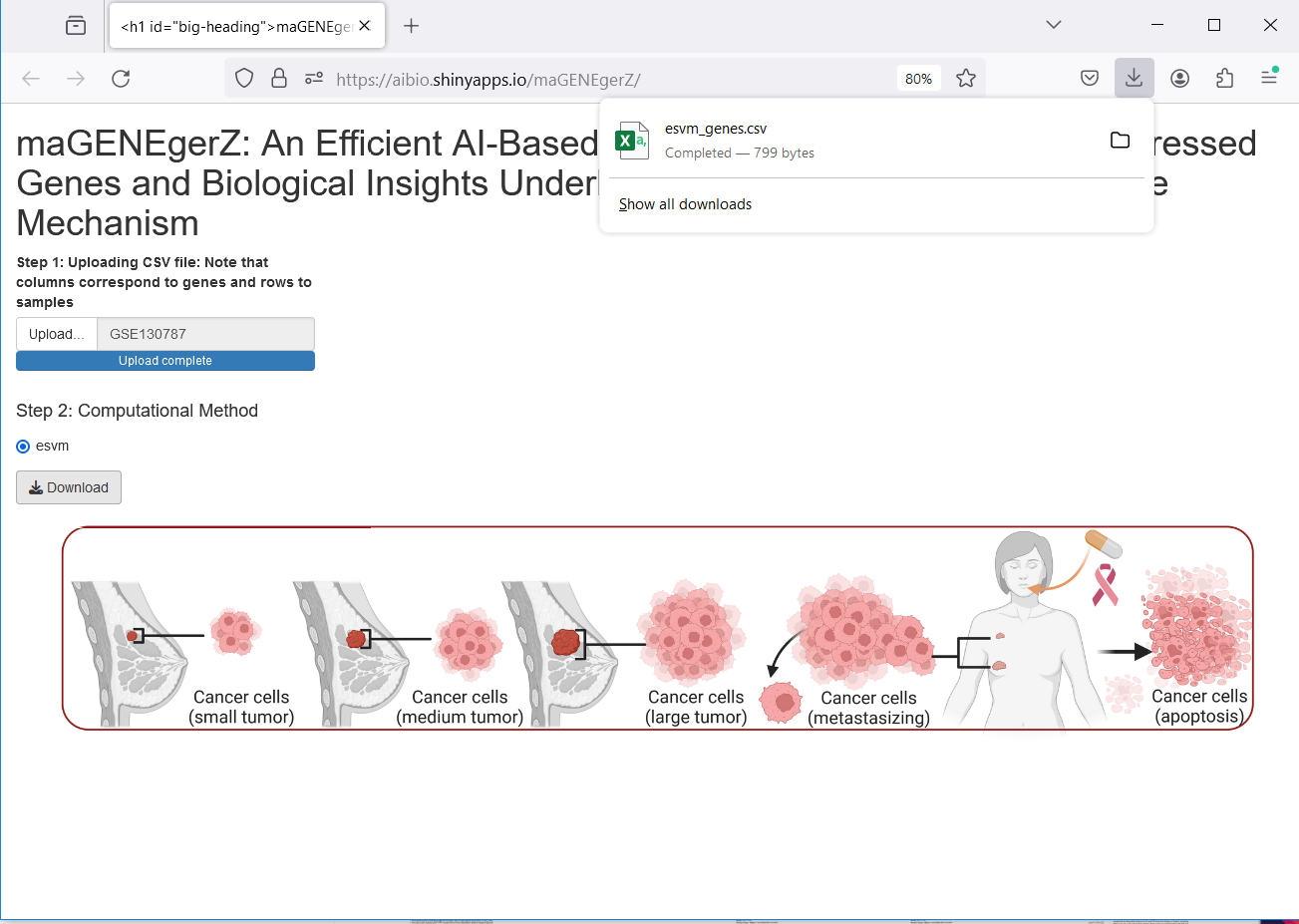

-Now, we open the results in excel file shown as follows

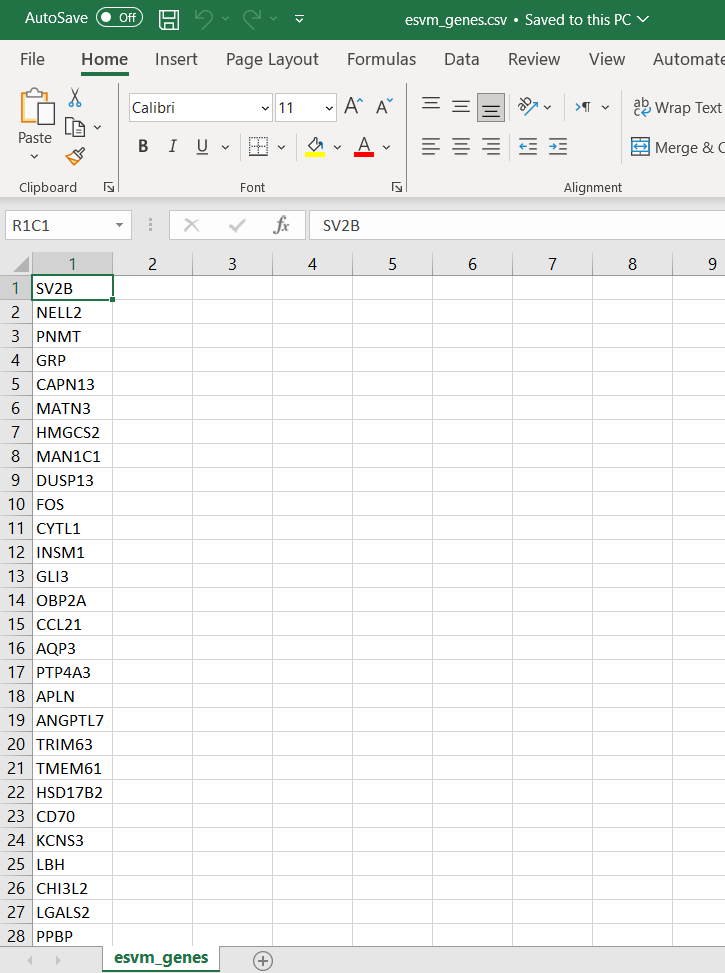
